# A Thermodynamic Model of Mesoscale Neural Field Dynamics: Derivation and Linear Analysis

**DOI:** 10.1101/2020.06.25.172288

**Authors:** Y. Qin, A.P. Maurer, A. Sheremet

**Affiliations:** Engineering School of Sustainable Infrastructure and Environment, University of Florida, Gainesville, FL. 32611.; McKnight Brain Institute, Department of Neuroscience, University of Florida, Gainesville, FL. 32610.; Department of Biomedical Engineering, University of Florida, Gainesville, FL. 32611.

## Abstract

Motivated by previous research suggesting that mesoscopic collective activity has the defining characteristics of a turbulent system, we postulate a thermodynamic model based on the fundamental assumption that the activity of a neuron is characterized by two distinct stages: a sub-threshold stage, described by the value of mean membrane potential, and a transitional stage, corresponding to the firing event. We therefore distinguish between two types of energy: the potential energy released during a spike, and the internal kinetic energy that triggers a spike. Formalizing these assumptions produces a system of integro-differential equations that generalizes existing models [Wilson and Cowan, 1973, Amari, 1977], with the advantage of providing explicit equations for the evolution of state variables. The linear analysis of the system shows that it supports single- or triple-point equilibria, with the refractoriness property playing a crucial role in the generation of oscillatory behavior. In single-type (excitatory) systems this derives from the natural refractory state of a neuron, producing “refractory oscillations” with periods on the order of the neuron refractory period. In dual-type systems, the inhibitory component can provide this functionality even if neuron refractory period is ignored, supporting mesoscopic-scale oscillations at much lower activity levels. Assuming that the model has any relevance for the interpretation of LFP measurements, it provides insight into mesocale dynamics. As an external forcing, theta may play a major role in modulating key parameters of the system: internal energy and excitability (refractoriness) levels, and thus in maintaining equilibrium states, and providing the increased activity necessary to sustain mesoscopic collective action. Linear analysis suggest that gamma oscillations are associated with the theta trough because it corresponds to higher levels of forced activity that decreases the stability of the equilibrium state, facilitating mesoscopic oscillations.

## 1. Introduction

A persistent challenge in understanding the neurobiological basis of higher-cognition is uncovering the mechanism by which neural activity across different scales of the brain is coordinated [Allen and Collins, 2013, Lashley, 1958]. At cell scale, action potentials (~10^3^ Hz) provide the “atomic” constituents of activity [Buzsáki, 2006, Buzsáki and Draguhn, 2004, Eichenbaum, 2017, Hasselmo, 2015, McNaughton et al., 1996]. At global-brain scale, the large-amplitude theta rhythm, with a frequency three orders of magnitude lower (6-9 Hz), is believed to provide a temporal structure around which smaller scale oscillations organize [Buzsáki, 2002, Green and Arduini, 1954, Green and Machne, 1955, Lisman and Idiart, 1995, Vanderwolf, 1969]. However, neither spikes nor theta in isolation can represent cognition, which suggests that neural dynamics fundamental for higher cognition reside in collective activity occupying a scale intermediate (meso-) between theta and action potentials. Following previous work [e.g., Freeman, 2000b, Muller et al., 2018b] we define here the mesoscale as spanning temporal scales between, say, 8 ms and 20 ms (e.g., LFP oscillations between 50 Hz and 120 Hz), and spatial scales in the order of mm to cm). These intervals correspond to the gamma activity [Bragin et al., 1995], prominent in the hippocampus.

At mesoscopic scales, the spatial organization of neurons within a neocortex layer shows a relative homogeneity. The mesoscopic neural activity supported by these layers involves a large number (e.g., ∼ 10^4^ − 10^8^, e.g., Deco et al., 2008) of synchronized action potentials that assemble into spatio.temporal patterns [Hebb, 1949, Lashley et al., 1951]. Should neurons be organized in a manner that favors local connectivity over long-distance projections, the spatio-temporal pattern of activity may manifest as propagating waves [Lubenov and Siapas, 2009, Patel et al., 2012, 2013, Petsche and Stumpf, 1960, Muller et al., 2018b]. Recent studies correlating hippocampal LFP to active exploration shows that neural activity develops as perturbations, spanning a wide frequency range, of a largely scale-free (∝ *f*^−*α*^) background state [Sheremet et al., 2016b, 2019b]. Following Freeman [2000a,b], we will refer to these perturbative patterns of neural activity as “mesoscopic collective activity”^2^.

The nonlinear, stochastic character of mesoscopic collective action suggests that the turbulence theory might provide an adequate framework for studying mesoscopic activity dynamics [Sheremet et al., 2019b]. In broad terms, turbulence may be described as a theory of the internal energy balance in nonlinear, systems with a large number of components whose dynamics spans a wide a continuum of scales. Nonlinearity implies interaction across scales, allowing for a cross-scale flux of energy. In cases where the cross-scale flux has a dominant, well-defined direction, it is often called “turbulent cascade” (e.g., figure 1). Turbulence was originally formulated as a general hydrodynamic theory, but has evolved to become the theoretical foundation of disciplines ranging from plasma physics, nonlinear optics, Bose-Einstein condensation, water waves, aggregation-fragmentation processes, and many others [Kolmogorov, 1941, Richardson, 1922, Zakharov et al., 1992a, Frisch, 1995, Nazarenko, 2011]. A key finding of the weak turbulence theory is the existence of equilibrium states of the multi-scale system, characterized by a self-similar distribution of energy across scales (the Kolmogorov-Zakharov spectra, Zakharov et al. 1992a, Zakharov 1999). In the research into brain activity, a concept that has some similarities is the “self organized criticality” hypothesis [e.g., Bak et al., 1988, Beggs and Plenz, 2003].

**Figure 1.**
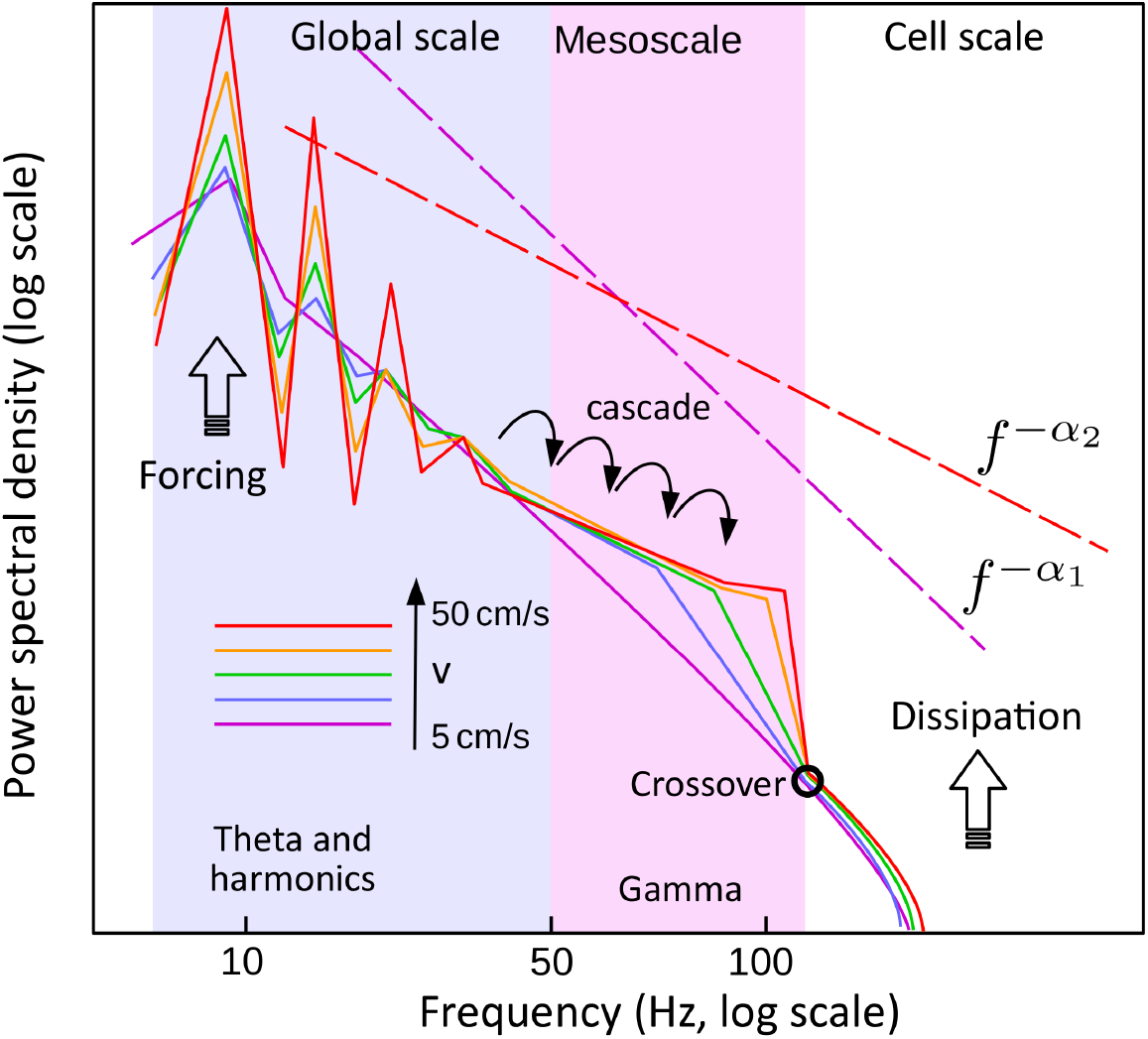
A cartoon of the typical evolution of the power spectrum of the hippocampal LFP with rat speed, that summarizes observations discussed in Sheremet et al. [2019b]. The evolution of the spectrum shows remarkable ordering by speed (e.g., from 5 cm/s to 50 cm/s, violet to red). Power increases by a factor of 4 in the theta band (blue rectangle), with theta and harmonics becoming prominent, while the gamma band exhibits a transformation that could be described as a spectral front shifting toward higher frequencies, up to the upper bound of the gamma band (black circle, crossover point), beyond which the spectrum no longer responds to forcing. This evolution suggests that nonlinear interactions between different frequency components result in a behavior similar to a turbulent cascade: the power received from external forcing in the theta band generates a net spectral power flux from low frequencies (theta) toward high frequencies. The crossover point (black circle at about 130 Hz) signals a significant shift the dominant physics. On the left side of it, in the gamma frequency band, nonlinear interactions dominate; on the right side physics are dominated by dissipation. The fundamental difference between the gamma activity and higher-frequency (cell-scale) activity supports the hypothesis that collective activity is macroscopic with respect to cell-scale processes. The spectral evolution is associated with a change in the overall slope of the spectrum (*α*_1_ corresponds to low speeds; *α*_2_ to high speeds).

Because mesoscopic collective action is macroscopic with respect to cell scale processes, previous research into mesoscopic brain activity has approached the problem either using the statistical-physics formalism [e.g., Nykamp and Tranchina, 2000, Cai et al., 2004, Ly and Tranchina, 2007, Rangan et al., 2008, Bressloff, 2011] or thermodynamics/hydrodynamic formulations (e.g., Wilson-Cowan class of fundamental equations, Wilson and Cowan, 1972b, 1973, Cowan et al., 2016, Amari, 1975, 1977, Deco et al., 2008) The statistical-physics approach characterizes macroscopic states by probability densities (configurations) of microscopic stats, and derives macroscopic equations applying averaging operators to microscopic physics. The thermodynamic approach defines the macroscopic state in terms of observable (macroscopic) state variables and postulates their balance equations. The statistical description founded on microscopic dynamics. It can capture in principle the full statistical details; in practice, however, it inherits from microscopic dynamics a very large number of degrees of freedom. The resulting equations may be very complicated and give rise to closure problems. The thermodynamic approach is simpler, effective, and is easy to construct, but at least in principle in principle to more limited than the statistical physics approach, due to fundamental quasiequilibrium assumption and its postulated foundation.

This study is motivated by long-term goal of understanding mesoscopic collective activity in the framework of the turbulence theory. Here, we introduce a new thermodynamic formulation of mesoscopic collective activity, and discuss its basic linear properties.

We adopt the thermodynamic formulation, both because its relative simplicity and its well-established history. The key equations were derived by Wilson and Cowan [1972b, 1973] and further refined by Amari, 1975, 1977, Wright and Liley, 1995b, Jirsa and Haken, 1996, 1997, Robinson et al., 1997, Cowan et al., 2016 and many others (see, e.g., reviews by Deco et al. e.g., 2008, Coombes et al. e.g., 2014, Cowan et al. e.g., 2016; because of their common fundamental principles, we refer below to models that are based on the Wilson-Cowan and Amari formalism as WC/A models). The model presented here, which belongs firmly to the WC/A class of models, was derived in response to the realization that all models of this class contain a curious deficiency. While the deficiency not detract from the value and success of the WC/A models, it does make current formulations ill suited for investigating turbulent aspects of mesoscopic brain activity. Indeed, the Wilson-Cowan (WC) class of models generally are formulated as a relationship between the local firing rate and incoming pulses in the element of area. In thermodynamics, this is largely equivalent to describing the evolution of a physical system only in terms of its exchanges with the external systems, i.e., in term of process variable. Because no state variables are defined, therefore the state of the system remains unknown. Amari’s [1975, 1977] approach corrected the issue to a degree, however, one may argue that the use of an “averaged membrane potential” as state variable may lead to difficulties because the quantity is ill defined during the explosive depolarization of a spike (Amari did not, in fact elaborate on the definition of this quantity). However, an explicit and accurate characterization of the state of the system is essential for investigating a turbulent system, because the distribution of the state variable over the internal scales of the system is related to the distribution of energy, which drives the energy cascade, i.e., the evolution of the system itself.

It is possible that this deficiency is the result of an original lack of interest in a rigid thermodynamic formalism, maybe too fastidious for many practical purposes. While correcting this deficiency is in itself a relatively small point, a consistent thermodynamic formalism has, however, a number of advantages: it provides a clear statement about the physical postulates underpinning the model; it defines state and process variables; it allows for an explicit description of the energy redistribution over scale in the collective activity system. The process also requires some changes in the formulation of standard functions such as the activation function. The resulting model is different enough from its “parent” WC/A class to warrant a closer examination of its basic properties.

section 2 discusses LFP measurements that form the basis of the turbulence hypothesis. We provide a short review of the WC/A class of models in section 3. In Section 4 we discuss what has arguably become the standard dynamical-kinetic-thermo/hydrodynamic modeling framework used for the representation of physical systems; we introduce the powder-keg paradigm, and we derive the governing equations of the thermodynamic model. The powder-keg model is compared to the standard Wilson and Cowan [1972b, 1973] and Amari [1975, 1977] models in section 5. Elementary simplifications that bring the equations to an analytically-tractable form are discussed section 6, and some rudiments of linear analysis are presented in sections 7 and 8 single- and dual-type neural fields. We conclude with a discussion of the results (Section 9). Details of the formulation of the new activation function, the positive-definite character of the state variables (internal “kinetic” energy and excitability), and algebraic details of the growth rate and dispersion relation derivation for dual-type neural fields, are given in the appendices.

## 2. Motivation

Recent investigations of hippocampal LFP in rats show a strong relation between energy input into the hippocampus (as inferred based on rat speed) and the nonlinear character of neural activity [Sheremet et al., 2016b,a, 2019b,a]. Both spectra and bispectra are well ordered with input power, as parameterized by rat speed. The redistribution of increased power over scales (frequencies) shows remarkable organization, as sketched in figure 1. In summary:

- At low frequencies, the power increase is highly localized to theta and its harmonics. Theta power increases by a factor of 4 and becomes strongly nonlinear (highly skewed and asymmetric; up to 5 harmonics can be clearly identified, Sheremet et al., 2016b). Frequency bands adjacent to theta and harmonics (e.g., *f* < 6 Hz, or 10 < *f* < 14) show a marked depletion of power.
- At high frequencies, gamma power increases by a factor of 2, but its power increase distributes through a process that may be described as a front moving across scales: gamma modes grow and plateau sequentially, starting at the lower frequencies (*f* ≃ 60 Hz) and progressing toward higher frequencies.
- As power grows, gamma develops significant nonlinear coupling with theta.
- The process of redistribution of power over scales process is reversible: if power levels retreat to initial values, the initial scale-distribution of power (spectrum) is recovered.
- At the lowest levels of power observable, the scale-distribution of power is nearly self-similar (power spectrum of the form *f*^−*α*^, with *α* > 0). We refer to this as the background spectrum (state). The background spectrum may be identified with a dynamic equilibrium point, i.e., a state that may be maintained indeterminately, but requires energy input.

If one identifies mesoscopic collective action with the gamma band, our observations suggest that these processes are perturbations of a dynamical equilibrium state (background state), and that increased power input in the theta band triggers a scale redistribution of gamma power. This evolution is tantalizingly similar to the energy cascade in a turbulent system.

## 3. Short review of neural population models

The beginning of the development of neural population modeling can be traced back to Beurle and Matthews [1956], who proposed an “update” equation to describe the propagation of large scale brain activity in networks composed of excitatory neurons, with applications to problems ranging from understanding the generation of LFP rhythms to visual hallucinations [Nunez, 1974, Milton et al., 1993, Ermentrout, 1998, Larter et al., 1999, Curtu and Ermentrout, 2001, Robinson, 2006, Pinto and Ermentrout, 2001a, Amari, 1977, Freeman, 1975b, Huang et al., 2004, Deco et al., 2009, Coombes et al., 2014, Muller et al., 2014], and with approaches ranging from detailed descriptions of randomly connected neurons transmitting all-or-nothing signals to hierarchically structured networks whose dynamics involve multiply spatial and temporal scales [Amari, 1975, Jirsa and Haken, 1997, Robinson et al., 2002, Breakspear et al., 2003, 2004, Breakspear and Stam, 2005, Nunez and Srinivasan, 2006, Deco et al., 2008, Mejias et al., 2016, Breakspear, 2017].

The mass action description [Wilson and Cowan, 1972b, Da Silva et al., 1974, Jansen and Rit, 1995, Marreiros et al., 2008] may be the simplest approach to population modeling, actively used since the 1970s to understand LFP rhythms, and deriving naturally from the concept of activity synchronization (e.g., Kuramoto, 1975, Strogatz, 2000). The key assumption is that at some local scale the activity of individual neurons is strongly synchronized and coherent, and thus one may describe it as the mean activity of the local neural mass, with interacting “masses” of neurons, such as excitatory and inhibitory neurons in different layers of cortex, modeled by a small number of equations, each describing the mean activity of a distinct neural “mass”. Theoretical treatments with empirical synaptic and input-response functions are possible (e.g., Freeman, 1979, Jansen and Rit, 1995, Miller et al., 2003, Stefanescu, 2011, Jirsa, 2011). The “mass” approach provides the building blocks for brain network models (e.g., Freeman 1975b, Breakspear et al. 2004, Breakspear and Stam 2005, Wong 2006, Honey et al. 2007, Deco et al. 2009, Jirsa et al. 2010, Woolrich and Stephan 2013), which treat the cortex as a discrete network of dynamical nodes (the neural “masses”) coupled through the connectome, essentially incorporating neural “masses” into a larger system that helps to understand topological significance of connections in organizing cognition, and functional correlations across brain regions. It should be clear, however, that this approach is in its essence a large scale model that lumps laminar neuronal tissues into discrete mass points, and thus does not resolve smaller-scale details such as mesoscale spatio-temporal patterns. The approach is not universally accepted and may lead to contradictory conclusions regarding large-scale brain dynamics [Breakspear, 2017]. Neural “mass” models may be developed into more complicated representations. For example, instead of using the spatial mean, the state of the neural population, one could follow a statistical mechanics approach and describe the neural “mass” using the probability distribution of neuron states. Under the assumption that the diffusion approximation holds true, one may derive Fokker-Plank-type stochastic equations (e.g., Kardar, 2007b,a; for applications to neural masses see e.g., Friston, 2010, Omurtag et al., 2000, Fourcaud and Brunel, 2002, Harrison et al., 2005, Ma et al., 2006, Deco et al., 2008, El Boustani and Destexhe, 2009; or fractional versions, Linkenkaer-Hansen et al., 2001, Lundstrom et al., 2008), useful for describing the evolution of network synchrony.

A next step toward a more flexible description of collective neural activity is to discard the concept of a “mass” of synchronized neurons and treat the cortex as a continuum, with the properties of the local neural population changing continuously in space and time. This class of models are referred to as neural-field models (see e.g., Ermentrout, 1998, Coombes, 2003, Deco et al., 2008, Cowan et al., 2016, Breakspear, 2017, Muller et al., 2018b, as well Gerstner et al., 2014, Coombes et al., 2014, Troy, 2008, Hoyle and Hoyle., 2006, Winfree, 2001).Their distinguishing characteristic is the elimination of the “individual neuron” concept. Instead, the dynamics of collective neural activity is described by a small number of fields, say *φ*_*j*_(*x*, *t*), where *φ*_*j*_, with *j* = 1, ⋯, *N* are *N* variables that characterize completely (in the sense of closing the system of equations) the neural field. The first such model was introduced by Beurle and Matthews [1956], who proposed an “update” equation to describe the propagation of large scale brain activity in networks composed of excitatory neurons. The model was revisited and extended by Wilson and Cowan [1972a, 1973], Nunez [1974], and Amari [1977]. A major limitation of the early Beurle and Matthews [1956] neural field model was its neglect of refractoriness or any process to mimic the metabolic restrictions placed on maintaining repetitive activity. Wilson and Cowan [1972a, 1973] The landmark model of Wilson and Cowan [1972a, 1973] coupled excitatory and inhibitory populations corrected this issue, and was successfully used to understand pattern dynamics such as oscillations and hysteresis, that shed light on real biology. The model proposed by Nunez [1974] links synaptic action to action potential firings, which allowed for periodic-wave solutions and sustained oscillations. The novelty of the model proposed by Amari [1977] was the inclusion of the “average membrane potential” as a state variable, coupled with firing rate. By assuming Heaviside activation function, Amari successfully derived solitary wave solutions for the model which opened a world of theoretical approximation on integral type of neural field equations.

Toward the beginning of this century, field models gained increasing popularity, which brought increased, systematic scrutiny of their properties, and additional refinements. Ermentrout and McLeod [1993], Ermentrout [1998], Osan and Ermentrout [2001] proposed a model that introduced a state variable similar to the membrane potential in Amari’s model, by integrating the firing rate (incoming energy flux, a process variable), and conducted an analysis of the existence and stability of solutions, including wave fronts and traveling pulses. Jirsa and Haken [1996, 1997] modified the Wilson and Cowan [1972a, 1973] models to account for axonal-delay effects proportional to the span of connections, and thus allowed wave solutions that arise as result of axonal propagation. Interested in electrocortical waves, Wright et al. [1994], Wright and Liley [1995a, 1996], Robinson et al. [1997], Freeman [1991] introduced another population model of coupled excitatory and inhibitory neurons following earlier work by Freeman [1991], Their model could be in fact regarded as a variant of the modified the Wilson and Cowan [1972a, 1973] model accounting for axonal delay (similar to [Jirsa and Haken, 1996, 1997]), but including no refractory period, and with a specific temporal weighting function comprising effect of synaptic delay and depolarization decay.

Wave propagation, and in general, the evolution of spatio-temporal patters in the cortex, arguably plays a central role in understanding collective activity dynamics. One of the earliest systematical derivations of traveling wave front solutions (arguably a simplest wave-like pattern) is due to Ermentrout and McLeod [1993], Ermentrout [1998]; although derived in a highly restricted formulation, their results, such as estimated velocity of activity propagation, shed light on biological information transfer. The role of inhibitory neurons in the formation and propagation of collective activity waves in a neural field is one of the fundamental results of recent studies (although the mechanism is not fully understood; see e.g., Wulff et al. [2009], Castro and Aguiar [2012], Stark et al. [2013], Amilhon et al. [2015], Neske et al. [2015], Hattori et al. [2017]). The interactions between excitatory and inhibitory neurons are believed to play an essential role in the dynamics and information processing of neural populations. The Wilson and Cowan [1972a, 1973] model and derivatives and known to have a rich set of spatio-temporal patterns, including oscillatory solutions in dual-type networks (including excitatory and inhibitory neurons; Wilson and Cowan, 1972a, 1973, Nunez, 1974, Larter et al., 1999, Robinson et al., 2002, Breakspear et al., 2003, Robinson, 2006); traveling wave fronts [Amari, 1977, Ermentrout, 1998, Pinto and Ermentrout, 2001a]; periodic progressive waves [Nunez, 1974, Amari, 1977, Robinson et al., 1997]; standing pulse solutions [Ermentrout, 1998, Amari, 1977, Pinto and Ermentrout, 2001a]; spiral waves [Milton et al., 1993, Osan and Ermentrout, 2001, Huang et al., 2004]; and maybe others. These patterns have formed the basis for experimental observations regarding the generation of sustained and propagating activity patterns in several brain regions Pinto and Ermentrout [2001a], Ermentrout and Kleinfeld [2001], Wu et al. [2008], Muller et al. [2018a]. It is important to note that excitatory-inhibitory neuron interaction is not the only mechanism of pattern formation. Purely excitatory networks support oscillatory solutions and traveling pulses [Curtu and Ermentrout, 2001, Pinto and Ermentrout, 2001b] as well as periodic traveling waves [Meijer and Coombes, 2014]. While inhibition (inhibitory neurons, spike frequency adaptation and and refractoriness; Ermentrout and McLeod, 1993, Pinto and Ermentrout, 2001a, Huang et al., 2004) plays an essential role in the formation and propagation of these patterns, its source is not well understood: models tend to produce patterns that agree qualitatively with observations, but with large quantitative deviations from observations that are still unexplained. Curtu and Ermentrout [2001] showed that the ratio of absolute refractory period over time constant should be >5, resulting a oscillatory period derived is between 1.4 and 4 in refractory period units. Likewise, propagating pulses and periodic waves discussed in the works of Pinto and Ermentrout [2001b] and Meijer and Coombes [2014] have time scales of the same order of magnitude as absolute refractory periods, which does not agree with large ratio of absolute refractory period to membrane reaction time necessary for sustained propagating patterns (in the Wilson-Cowan model the absolute refractory period needed for propagating waves is in the order of 10 time-constant units (at least 40 ms, while membrane reaction time, or time constant, is ≈ 10 ms).

While this brief review of collective activity models does not even come close to doing full justice to all the research effort dedicated to the problem, it should highlight some of the peculiarities of its history: the brilliant and rather ad-hoc ideas, the late intersection of their evolution with other well-developed, mature branches of physics such thermodynamics, statistical mechanics, and kinetics. This is reflected in the peculiar usage of state and process variables, the lack of a systematic approach to the study of the dynamics of spatio-temporal patterns. Interestingly, this is not for the lack of enthusiasm (e.g., Freeman, 2000a,b, 2006, 1975a, Freeman and Vitiello, 2010, 2006 to cite one of the most enthusiastic investigator of collective activity). Still, the remarkable persistence of the Wilson and Cowan [1972a, 1973] model as a key, fundamental formulation for neural-field activity is reflected in that all subsequent models are closely related to the original delayed form of Wilson and Cowan [1972a, 1973] equation, either directly deriving from it, or reduce to it through time coarse-graining. This implies that the mechanisms and capability of field models have changed little over a long history, and suggests that their rich reservoir of solutions met most expectations in terms of reproducing occasionally observed patterns in recordings. This may also, however, be the result of rather intermittent, occasional interest in collective activity (stemming mostly from practical computation interests), perhaps obscured by the dominance of the philosophical view known as “multiplexing”, that postulates that neurons function in a way similar to electronic components hardwired on a circuit board in a computer. If the latter were true, then collective activity would be indeed at most of a secondary concern. However, as observations and hypotheses accumulate that contradict the “multiplexing” model, such as the degeneracy and role of turbulence and self-organized criticality in collective neural activity (e.g., Edelman, 1987, Edelman and Gally, 2001, Beggs and Plenz, 2003, Shew et al., 2011, Beggs and Timma, 2012, Sheremet et al., 2018a, 2019b and others), or perhaps simply due to the growing interest in mesoscale processes, the capabilities of the Wilson and Cowan [1972a, 1973] model are bound to undergo further scrutiny.

So far, collective activity patterns have been studied from a perspective reminiscent of the theory of pattern formation in dynamical systems, in the sense that particular patters have been identified and studied in isolation. Solutions in a given model with physiological parameters determined are confined by and large to a single scale. However, waves generated in a single brain region are never confined in a single scale, but always corresponds to a spectrum spanning at least the domain from 1 Hz – 300 Hz. The dynamics of the spectral distribution of energy in a hippocampus LFP raises a number of questions (e.g., Sheremet et al., 2017, 2018a, 2019b) that cannot be addressed directly using the current formulations. While the value of the Wilson and Cowan [1972a, 1973] formulation is beyond dispute, a number of small changes are needed to address the problem of the spectral evolution. The rest of this paper is dedicated to the discussion of these modifications.

## 4. A thermodynamic mesoscopic model for neural fields: the powder-keg paradigm

### 4.1. Microscopic vs macroscopic^3^

The words “macroscopic” and “microscopic” are used here as a non-dissociable pair of relative terms, that define two fundamental scales coexisting in the system, governed by fundamentally different physical laws. The microscopic scale refers to processes that involve some atomic (in the etymological sense of “not further divisible”) elements of the system. If the system has a large-enough number of atomic elements, collective behavior might emerge, in which the contributions of individual atom are indistinguishable (e.g., atoms may conceptually be interchanged without altering the collective behavior). Such processes are macroscopic, and are governed by physical laws effectively different that atom-scale processes^4^. The definition of the dual micro/macro scales is arbitrary, determined by the processes of interest. Micro- and macrodynamics coexist: for example, while individuals participating in a stadium wave may eat, read a newspaper, chat in pairs, etc, to create a stadium wave all they are asked to do is stand and sit in synchrony with the rest of the group.

The word “scale” is used below with two additional meanings. As common in physics, the generic term “scales” is used to refer to wave numbers or frequencies in the Fourier representation. Neuroscience also defines two absolute scales: the “brain (or global) scale”, and the “cell scale”. The global scale refers to processes that span a significant part of the entire brain. The cell scale refers to processes that involve individual neurons, the natural “atoms” of the cortex, whose physics are described, say, by the Hodgkin and Huxley [1952] model.

Therefore, ignoring sub-cell processes, we will define here the cell-scale as microscopic. The definition of the dual macroscopic scale deserves more discussion. Following the reasoning discussed above, the macroscopic scale is the scale where collective behavior emerges. The existence of a spectral crossover point in the neighborhood of 130 Hz (figure 1), suggests that gamma oscillations are governed by different physics than the cell (microscopic) scale, rons entrained in these processes, say, in the order of *O* (10^4^) seems small, it is important to implying that mesoscopic activity is macroscopic in relation to cell scale. If the number of neunote that order of magnitude of the number of atomic constituents needed for macroscopic behavior is not an a priori given number, but depends on the system under consideration. moreover, the emergence of macroscopic behavior also depends on microscopic mixing, i.e., strength of interaction between atomic components. Strong microscopic mixing promotes macroscopic behavior. In his sense, micro-macro duality is the expression of the dynamics of the system, and not of some absolute number of components. This observation has important consequences for brain activity. If neurons are hardwired like fixed electronic circuits, in unique patterns that assign neurons unique specific functions, there can be no mixing, no microscopic randomization, and therefore the “macroscopic” behavior is trivial (and irrelevant). However, evidence suggests that this is not the case. Synapses have a limited life-span, lasting only a few weeks [Attardo et al., 2015, Holtmaat et al., 2005, Xu et al., 2009, Xiao et al., 2009]. Mossy fibers from a granule neuron have up to 200 different synaptic inputs onto a wide variety of neurons [Amaral et al., 2007] and a single pyramidal neuron has over 30,000 synaptic inputs (e.g., Megias et al., 2001). These observations indicate that circuit model descriptions are not suitable for the cortex Maley [2018], and that the cortex structure is consistent strong nonlinear mixing [Buzsaki, 2006] and degeneracy [Edelman, 1987, Edelman and Gally, 2001]. We hypothesize that mesoscopic processes are macroscopic with respect to cell scale.

The distinction between macroscopic and microscopic descriptions (models) is particularly useful for systems whose exact microscopic state is impossible to measure. Although macroscopic dynamics should arguably be the direct result of microscopic dynamics, an explicit and formal derivation of macroscopic laws starting from microscopic physics is in general extremely difficult to construct. There are only a handful of very simple physical systems for which this connection is well understood (e.g., Alexeev, 2004, Kardar, 2007b). For practical purposes such a derivation is also in general not needed (see also the discussion below).

### 4.2. Dynamical, kinetic, and thermodynamic/hydrodynamic descriptions

Historically, the dynamical, kinetic and hydrodynamic/thermodynamic approaches for modeling physical systems with a large number of components were developed to explain how macroscopic physics emerges from microscopic dynamics. Statistical mechanics and kinetic theory are well understood for particle systems, and have been later generalized to other fields (e.g., magnetization) with various degrees of detail. The ideas below are elementary and may be found in any textbook of statistical mechanics textbook, (e.g., Gibbs 1902, Tolman 1938, Khinchin 1949, Kittel 1958, Pathria and Beale 2011 and many others) and kinetic theory (e.g., Boltzmann, 1872, 2003, Alexeev, 2004, Pathria and Beale, 2011, Kardar, 2007b,a, Tong, 2012 and many others).

Because the goal of this study is to formulate a thermodynamic model of collective (mesoscopic) activity, we provide here a sketch of these stages of modeling. Consistent with the fundamental work of Wilson, Cowan, and Amari, we follow what we believe is a consistent line of reasoning that allows for formulating the macroscopic laws governing collective activity.

#### 4.2.1. The dynamical model

A dynamical model is the collection of the evolution equations that describe the dynamics of each microscopic atomic component. In the case of an ideal gas made of a large number of identical particles, the fundamental law of mechanics Arnold [1974] states states that the mechanical state of a particle is completely defined by 6 degrees of freedom (three position components and three velocity components). For the brain, the number of equations included in the dynamical description equals the number of neurons described, times the number of degrees of freedom that describe the cell in, for example, the Hodgkin and Huxley [1952] model. Note that the dynamical model is fundamentally “phenomenological”, i.e., assembled together based on its capability of describing what is assumed to be the most relevant features of cell dynamics (in this case, the action potential). Its privileged status of fundamental model comes for the decision to ignore the sub-atomic (i.e., subcell in the case of the brain) physics. The dynamical equations are “deterministic” in the sense that if the initial conditions were known for each molecule, the equations of motion could, at least in principle, be integrated exactly. However, for practical applications the dynamical system is largely useless, for at least two reasons: the system of equations is too large to be solved directly in any practical application, and the exact initial conditions are not known.

For simplicity, we postulate that all neurons of a given type (e.g., excitatory) are physiologically identical^5^. Because this prototypical neuron may be defined through some averaging, it will be referred to as “mean neuron”. We assume that the key dynamics of the mean neuron are described by the standard “leaky integrate and fire” model of the action potential (figure 2.a). For example, the mean neuron is excitable if its membrane polarization is subthreshold (≲ −50mV, state (A) in figure 2.a). In this state the average potential fluctuates approximately between −70mV and −50mV^6^, due to small post-synaptic potentials, ion currents associated with membrane channels, etc. In state, the neuron is “excitable”, i.e., ready to fire. We refer to this state as the “background” state. If synaptic input is zero, the potential of the mean neuron decays to the resting state (≃ −70mV). If the input stimulus is large enough (state B in figure 2.a), it can trigger a spike (state C). After the spike, the neuron enters the hyperpolarization stage and slowly depolarizes (state D), returning to the original mean state (A). State (B) may be seen as a perturbation of the mean state (A), that triggers firing. During the spike (C) the neuron is “unavailable”, it does not respond to stimuli (absolute refractory state). In the hyperpolarization/recovery stage (D) the neuron is in relative refractory state: it is excitable, but it requires more energy input, relative to the background state (A), to trigger an action potential.

**Figure 2.**
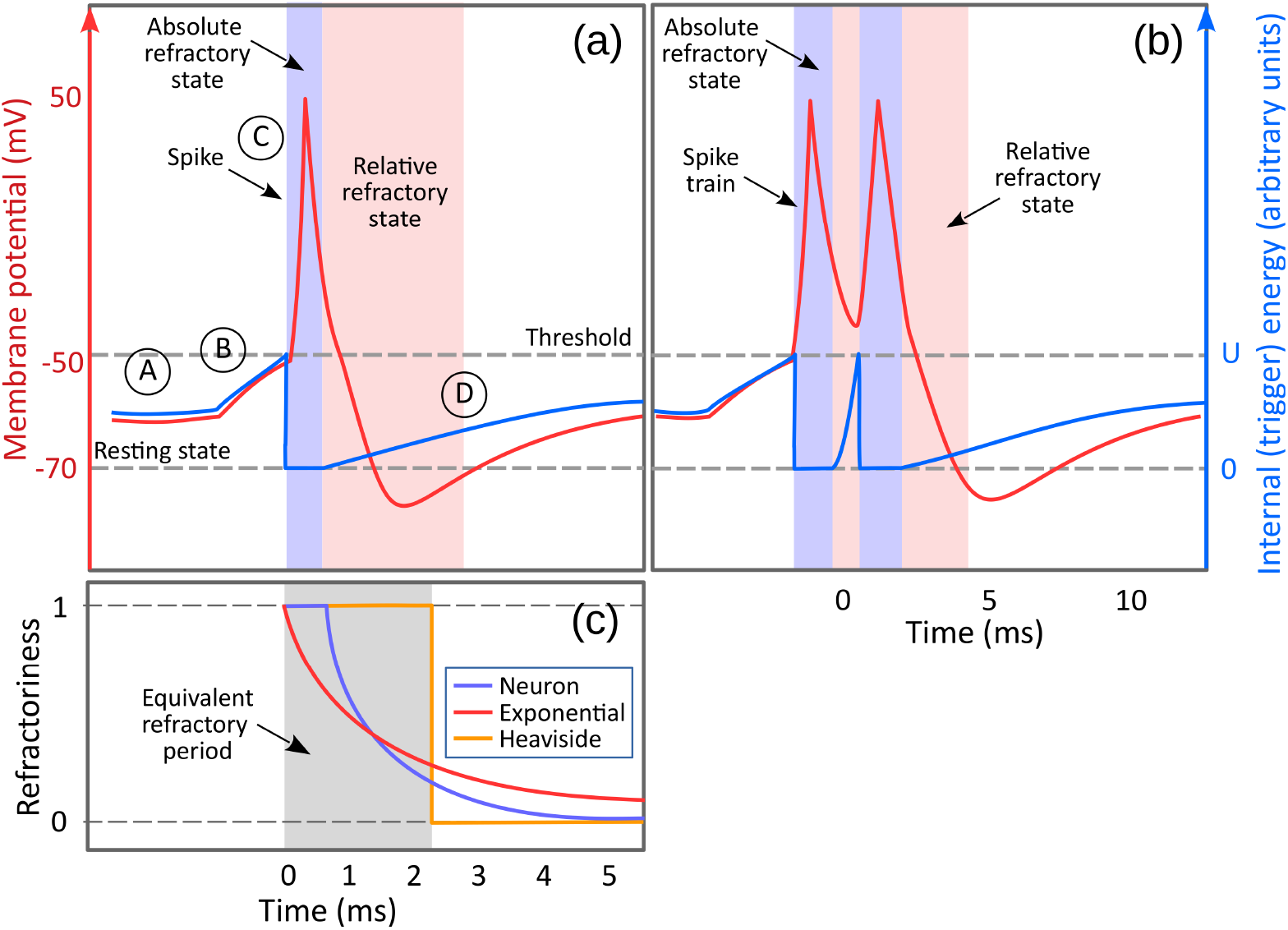
A cartoon of the standard “leaky integrate-and-fire” neuron model. *a)* Typical representation of the mean potential evolution including a potential spike (purple) and the definition of “trigger” kinetic energy (blue, see also text for a discussion of the meaning of “kinetic”). The evolution of membrane potential as a succession of states that may be described as “background” state (A); perturbations of the background state that bring the membrane potential to the threshold value (dashed gray), triggering a spike; action-potential spike (A); and post-spike state (D) in which the membrane potential goes into the hyperpolarization state and slowly depolarizes back to the background state (A). Refractory states are represented as colored vertical bands: in the absolute refractory state (blue) the neuron does not respond to stimuli; in relative refractory state (pink) the neuron is increasingly responsive, but the energy input required to fire is higher than in the background state (the excess input needed recedes as the neuron depolarizes). In reality, the average membrane potential is ill defined during the spike, therefore it cannot be used to describe the state of the neuron. The kinetic (trigger) energy of the neuron (blue line), is roughly proportional with the average membrane potential, it is bounded between zero (resting state) and the threshold value (*U*). As a thermodynamic quantity, the kinetic energy is defined in relation to the neural field, therefore it has no meaning when the neuron is not responsive to stimuli, therefore it is set to zero during the absolute refractory period. *b)* Stronger and longer-lasting stimuli may force the neuron into a spike train. Spike trains are represented here as rapid successions of single spikes. A single spike is produced by a short-lived perturbation (B) of the background state that brings the kinetic energy to the threshold and then disappears. If the perturbation is longer than a spike and strong enough, it can trigger a sequence of spikes in rapid succession. *c)* different representations of the refractoriness *r* of the mean neuron for a single spike and a spike train. The refractoriness *r* is a real number between 0 and 1 that reflects both the absolute and relative refractory states (see text for a discussion). The values of the membrane potential given here are for illustration purposes only; in actuality they depend on the type of neuron considered.

#### 4.2.2. The kinetic model

The kinetic theory is the first step toward a macroscopic description. The macroscopic state has by definition a much smaller number of dimensions, therefore one macroscopic state must correspond to a large number of microscopic configurations (e.g., Kardar, 2007b). Because the exact microscopic configuration is not accessible the macroscopic level, the macroscopic state of the system is described by *n*-component, joint probability density functions (PDF). A statistical description of the system amounts to a set of equations that describe the evolution of these distributions. The number of unknown functionals remains still dauntingly large, but some progress may be made is one restricts the effort to describing the PDF of a single component (e.g., macroscopic observations are local averaging operators based on the 1-component PDF). However, the evolution of 1-component distribution depends on the 2-particle distributions, which in turn depends on 3-particle one etc.

This hierarchy of dependencies is known as the BBGKY hierarchy (Bogoliubov-Born-Green-Kirkwood-Yvon e.g., Alexeev, 2004, Kardar, 2007a, Tong, 2012). The system of equations is infinite and unsolvable unless a closure exists. The celebrated Boltzmann equation, also known as the kinetic equation, is an example of quasi-Gaussian closure, where the higher-order joint PDFs may be factorized into products of the 1-particle PDF. The kinetic description is stochastic, in the sense that two macroscopic states (characterized by he same density function) represent many distinct realizations of microscopic configurations. This approach may be characterized as neither entirely microscopic nor entirely macroscopic: while the exact microscopic state is not specified, some information about the microscopic states is preserved in the probability density functions.

Accepting for now the conventional description of membrane-potential evolution shown figure 2, the kinetic state of the neural population is characterized by the PDF of membrane potential (figure 3). At any time *t* and position *x* the fraction of the neural population with the potential below the threshold is excitable in various degrees and may be triggered to spike; the rest of the population is firing (absolute refractory time). A fraction of the energy of the spike is passed along to other neurons through network connections; the rest is lost through various processes, such as electromagnetic radiation and ineffective connections. The background state could be interpreted as a steady, spatially uniform state in which the energy recaptured from spikes matches exactly the loss of internal energy to maintain its global mean energy level (dark green line in figure 3). In this representation, mesoscopic action processes are perturbations of the background state that locally change the membrane potential (bright green line). For example, a local increase in the internal energy shifts the distribution of neuron trigger energy toward the threshold, increasing the firing rate, and, as a consequence, the amount of recaptured energy and the internal energy of the system.

**Figure 3.**
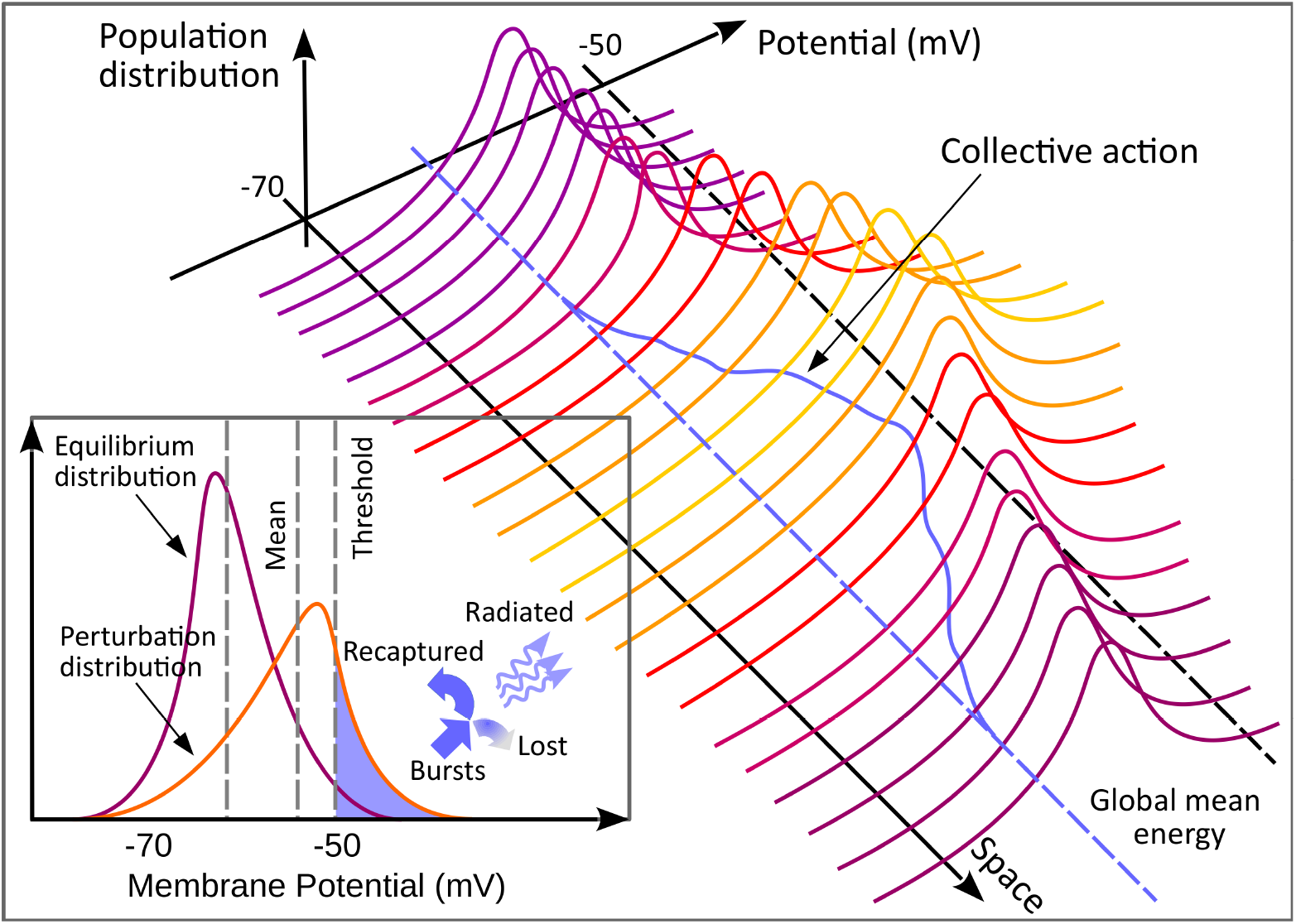
A cartoon of the probability distribution of the mean membrane potential over the neuron population in the element of volume. Although the mean membrane potential is ill defined during a spike (see figure 2), we use this representation for convenience. Using for this the correct state variable (trigger energy) is complicated due to being reset to zero after a spike. *Main panel*: Spatial structure of the distribution of internal energy over neuron population. The global mean energy is represented by a blue dashed line. The mean energy is represented by a continuous blue line. Continuous deviations of the mean energy from the global mean represent collective activity. *Inset:* Sketch of possible shapes of the distribution of average membrane potential over the neuron population for the equilibrium state (A) in figure 2.a (purple) and perturbed state (mesoscopic collective action, B in figure 2.a, yellow). The high value tails of both distributions exceed the threshold, implying that an number of neurons fire in both cases. The number of firing events is much larger for the perturbed distributions. In general, the shape of the distribution changes as the mean shifts, therefore the population exceeding the threshold (firing rate) depends not only on the mean but also on the distribution shape (higher moments). A fraction of energy released by spikes (blue arrows) is recaptured by the neural field, and the rest is lost to a host of physical processes.

#### 4.2.3. The thermodynamic limit

If the system is at macroscopic equilibrium or if its evolution is not too fast, the kinetic equation may be recast in the regular thermodynamic/hydrodynamic conservation laws (e.g., Alexeev, 2004, Tong, 2012). These equations are truly macroscopic, in the sense that all information about the existence of a microscopic structure is lost and replaced entirely with a macroscopic description. For example, the flow of fluid is completely described by the fields of pressure and flow velocity. This description is again deterministic: if the macroscopic state is known accurately, the future macroscopic state is exactly predictable. It is important to note that thermodynamic models have been (and still are) developed without the need of an explicit representation of, and derivation from, the underlying microscopic physics. This is in fact the whole point of the “macroscopic” concept: the governing laws are formulated for the observable (macroscopic) reality; the microscopic world is not observable. In this sense, any physical model is phenomenological.

The full modeling cycle starting from the dynamical description and ending in the thermodynamic limit has been examined is detail only for a handful of systems (e.g., Alexeev 2004). The vast majority of physics is based on phenomenological models whose connection to some underlying microscopic structure either is not well understood, or is inconsequential for the macroscopic description. In the brain duality of microscopic (cell-scale) to macroscopic (collective activity) scales, the WC/A class of models belong to the thermodynamic limit.

#### 4.2.4. Collective-activity turbulence

If collective activity is macroscopic with respect to cell scale, then the WC/A class of models (or generalizations, see below) should provide an adequate modeling platform for testing the mesoscopic turbulence hypothesis. The turbulence formalism is a field theory (e.g., Goldstein et al. 2014, Kardar 2007b,a, Tong 2012; many others) that describes the internal redistribution of energy (and other conserved quantities) over the Fourier scales spanned by the system. The equations governing both hydrodynamic [Richardson, 1922, Kolmogorov, 1941, Frisch, 1995] and wave turbulence [Zakharov et al., 1992a, Newell et al., 2001, Nazarenko, 2011] belong to the hydrodynamic class of equations, in the terminology discussed above. Applied to mesoscopic activity, turbulence describes the dynamics of multi-scale patterns of collective activity (not individual-cell activity). A brief introduction and references may be found in Sheremet et al. [2019b]. The Fourier components are the atomic components of the physical system. Because these components are macroscopic with respect to cell scale, WC/A models play the role of the dynamical model. The WC/A model could be solved directly for the evolution of each Fourier mode, but just as with the microscopic configurations of molecules in an ideal gas, we do not know the exact initial conditions (in this case, say, the initial phases). Spectral densities represent the distribution of power over patterns of different scales. This is a kinetic description, stochastic because the exact microscopic configurations (e.g., initial phases of the patterns) are not resolved. This description is implied in most of the data analysis techniques used to describe LFP characteristics; for example, the spectral density is an ensemble averaged quantity. A Boltzmann-type kinetic equation [Alexeev, 2004] may be derived following the blueprint of the BBGKY hierarchy and closure mechanism [Zakharov et al., 1992b, Nazarenko, 2011]. For gravity waves, this equation is known as the Hasselmann equation [Hasselmann, 1962]; for wave (weak) turbulence theory known as the Zakharov equation (Zakharov et al. 1992b, Zakharov 1999, Newell 2002, Nazarenko 2011 and others). One of the fundamental results of the wave-turbulence theory is the existence of self-similar spectra, called the Kolmogorov-Zakharov spectra. We hypothesize that this framework may help shed some light on the formation and the physical meaning of LFP spectra.

### 4.3. The powder-keg paradigm

The conventional representation of the action potential shown in figure 2 does not translate well into a quantity whose value can be used for thermodynamic purposes to describing the state of the neuron. The goal of such a state variable would be to characterize the state of a neuron as a whole by a single value, e.g., similar to the mean kinetic energy of a molecule in a gas. The mean membrane potential is a good candidate, because it is meaningful and descriptive for the microscopic sub-threshold equilibrium states, when the charge may be thought of as relatively uniformly distributed across the neuronal membrane. However, a spiking neuron is in a transitional (far from equilibrium) microscopic state, with charges highly localized as the electrical pulse propagates along membrane. In such a state, while a value for the mean membrane potential could still be defined, it is much less representative of the microscopic process.

Because a single-value characterization of the membrane potential of a neuron is not possible during a spike, for thermodynamic purposes, the sub-threshold state and the spiking process should be treated as two distinct processes. This suggests the thermodynamics of a powder keg. The term “powder keg” is used here to designate a thermodynamic device characterized by two distinct types of energy: a “potential”^7^ energy that is released as a spike (equivalent to the potential energy achieved by maintaining different concentrations of separation of sodium and potassium ions inside and outside the cell); and a “trigger” internal energy, proportional to the keg temperature (equivalent to the mean subthreshold membrane potential). When the temperature reaches a threshold level, it triggers the release of the potential energy (trigger voltage-gated ion channels). The powder keg thermodynamics is virtually identical to the conventional evolution of the membrane potential shown in figure 2: the background state of the neuron may be seen as the ambient temperature; when it reaches a certain threshold, the keg explodes, analogous to the neuron spike. The refractory state of a neuron may be simulated by replacing the exploded keg immediately after the explosion with an identical one whose temperature is initially zero and increases slowly to the ambient value through heat exchanges due to nearby explosions.

If the dynamics of the mean neuron is equated with that of a powder keg, a neural network may be represented as a large warehouse of powder kegs. The internal energy of the warehouse is defined as tho sum of the trigger energy of the kegs, and it is a variable independent of the potential energy released by explosions. Assume that the global mean temperature in the warehouse is somewhere between zero (no kegs explode) and the critical threshold temperature (all kegs explode). Local temperature fluctuations may cause spatially scattered explosions. A fraction of the energy released by explosions is recaptured by the system and increases the ambient temperature; the rest is lost to a variety of other processes such as light, sound, radiated heat, etc. In the absence of external energy input, explosions provide the only source of energy that can contribute to the ambient temperature. If no explosions occur, the temperature of the system naturally decays to a reference value (zero) below the threshold. An equilibrium state of the system is achieved if the energy recaptured from explosion balances the natural energy decay and other energy losses.

The distinction between internal (trigger) and potential energy in the powder-keg representation suggests adopting the simplifying assumption that the neuron spike (state C in figure 2.a) and non-firing states represent distinct processes, drawing from distinct pools of energy: 1) the potential energy released by a keg explosion, uses an accumulated source of energy, that is exhausted in a spike and needs to be replenished, and 2) the internal kinetic energy of the mean neuron, controlled by ambient network activity. The internal “kinetic” energy of the mean neuron is roughly proportional to average membrane potential (similar to Amari, 1975). We refer to this quantity as internal “kinetic energy” (as opposed to potential energy) because it is a direct expression of activity. For example, in an hypothetical “inactive” (but not dead) system, the mean neuron would be at resting state, i.e., its “kinetic” internal energy would be zero. Note that we adopt here the convention the internal kinetic energy is zero in the absolute refractory state, consistent with both the evolution of the neuron during the relative refractive state, and with the thermodynamic meaning of the internal kinetic energy as state variable of the system: if the neuron does not participate in the system dynamics, it does not contribute to the internal kinetic energy of the system. The powder-keg paradigm is a simplified thermodynamic (macroscopic) representation of a system of identical “leaky integrate-and-fire” neurons.

### 4.4. Governing equations

Below, the space and time are independent variables, measured in mesoscopic units, i.e., macroscopic with respect to cell scale. As a consequence, the duration of a spike is considered infinitesimal. If the neural field comprises several types of neurons, we denote the neuron type by superscript symbols, e.g., *E* for excitatory and *I* for inhibitory neurons. All neurons of a given type are assumed to have identical physiological properties.

Modeled in the powder-keg paradigm, the state of the neural field is completely defined by two independent state variables: 1) the internal kinetic energy ; and 2) the “excitability” of the neural population, that is the fraction of the population that is excitable. Because the energy exchange within the neural field is achieved through explosions (spikes and spike trains), the relevant process variable is a measure of the energy released by the fraction of the neuron population that is firing. In thermodynamics, state variables are extensive quantities. For convenience, we normalize here extensive variables by the number of neurons in the element of volume (intensive quantities).

The quantity *u* is defined as the internal kinetic energy in the element of volume at *x*^8^, normalized by the number of neurons. This is an intensive quantity that may be interpreted as the “temperature” (normalized kinetic energy) of the system (not to be confused with the standard temperature, measured by a thermometer; e.g., Callen, 1960). Therefore, 0 ≤ *u*(*x*, *t*) ≤ *U*(*U* is the threshold value). Neurons that have non-zero kinetic energy may be triggere and will be referred to as “excitable”.

Excitability is a property dual to refractoriness. The refractoriness of a mean neuron is measured by the fraction *r* of the incoming neurotransmitter flux that is ineffective, satisfying the conditions: *r* = 1 in absolute refractory state, 0 < *r* < 1 in relative refractory state, and *r* = 0 otherwise (e.g., figure 2.c). The dual parameter 1 − *r* may be used as a measure of “excitability” of a neuron. 1Let *N*(*x*, *t*) be the number of spikes per unit of time and volume, (it has the dimension of *t*^−1^)normalized by *ρ* (spike trains induced by strong and longer lasting stimuli are treated here as single spikes; e.g., figure 2.b). We will refer to this quantity simply as “firing rate”. Then, the refractoriness of the neural population may be written as 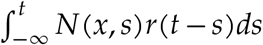, therefore the population excitability *a*(*x*, *t*), the fraction of neuron population not in the absolute refractory state, is

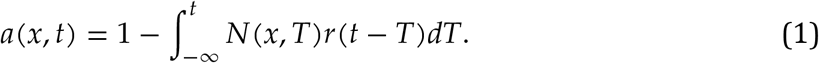

If the average energy captured by the neural field ε from a single spike is known, then *N* fully characterizes the internal kinetic energy exchange of the neural field.

Therefore, aside from variables that characterize the physiological properties of the network, the dynamical variables that describe the evolution of the neural field activity are the internal kinetic energy *u*(*x*, *t*), the population excitability *a*(*x*, *t*) (state variables), and the firing rate *N*(*x*, *t*) (process variable).

The processes governing the rate of change of the internal kinetic energy *u* are: 1) the incoming flux of depolarizing inputs coming through synapses; 2) the post-spike collapse of kinetic energy of the activated neurons, and 3) the natural tendency of the internal kinetic energy to decay due to microscopic dissipative processes (sodium-potassium ion pumping that restores the electrochemical gradient). The energy balance equation for the *α*-type neurons is therefore

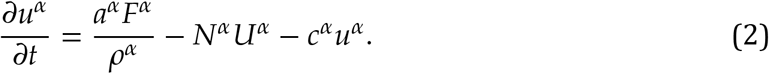

The first term in the right-hand side of equation 2 states that the contribution of the mean flux of energy *F*/*ρ* incoming through synapses to a neuron depends on the mean neuron excitability (e.g., if *a* = 1 all neurons *ρ* in the element of volume are excitable, the entire flux is absorbed). The input flux comes from connected neurons, and depends on the connection configuration, therefore it may be written in the general form

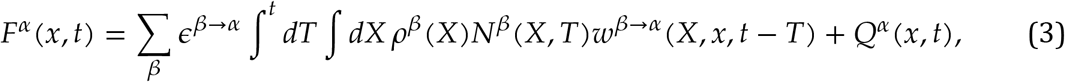

where ϵ^*β*→*α*^ is the average amount of energy released by a single spike from type-*β* neurons, as received by type-*α* neurons; *Q*^*α*^ is the energy flux arriving at *α*-type neurons; and *β* should be regarded as a variable that covers all neuron types, such that ∑_*β*_ is a summation over all types of neurons, including *α*-type ones. The spatial integration is carried over the spatial domain directly connected to the element of volume at *x*. The function *w*^*γ*→*α*^ is a weighting function that depends on the distribution of connections and axonal delays (see the appendix for the discussion).

The second term represents the post-spike loss of internal kinetic energy (figure 2). As discussed above, in the powder-keg paradigm the internal energy of a spiking neuron is set to zero, therefore, the process of releasing the potential energy *N* spikes annihilates *N*(*x*, *t*)*U* of the mean internal kinetic energy.

The third term describes the natural tendency of the kinetic energy to collapse to the zero-energy resting level in the absence of stimulus. Here, again we ignore the possible complexities of the decay-rate relation to mean energy, and assume that a constant decay rate *c* (perhaps to be refined at a later time) captures the essential character of the dynamics.

As discussed in more details in appendix A, we expect the probability of spiking to increase with higher depolarization (higher temperature). Therefore, the firing rate *N* should depend on the details of the internal-energy distribution over the neural population, i.e., on the mean internal kinetic energy *u* and higher moments therefore on all the moments of internal kinetic energy distribution), i.e.,

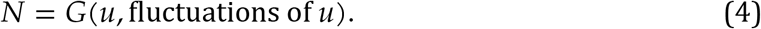

Because we are interested in this study in the leading order behavior of the system, in the absence of further guidance, and pending future refinements, we make the simplifying assumption that the distribution characterized primarily by its mean, and that the contributions of the fluctuations of the mean are not significant and may be neglected. Therefore, we replace for now equation 4 with the simple form

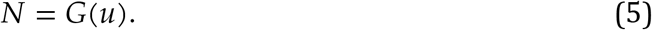

Collecting all above equations, one obtains the system

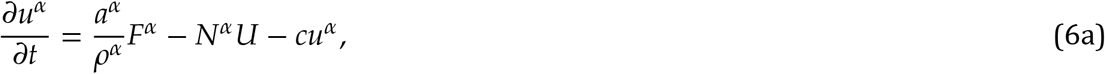

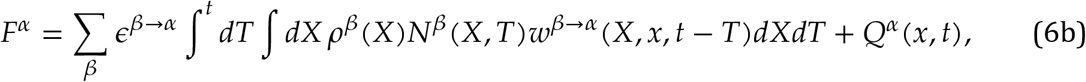

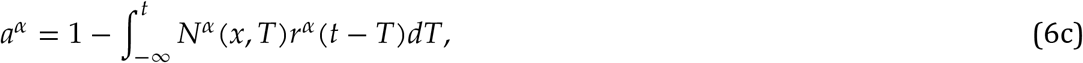

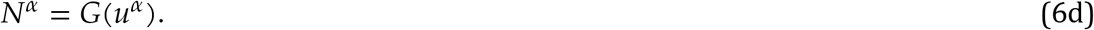

Equations 6 are the governing equations for the powder-keg thermodynamic paradigm of a neural-field continuum comprising several types of neurons. These equations are general, both in the sense that contain expression and parameters yet to be specified (e.g., equation 6d) and in the sense that described a wide variety of processes other than collective action defined as a perturbation of the background state. Equations 6 are complicated nonlinear integro-differential equations that are extremely difficult to interpret and solve in original form. They involve both a number of parameters (e.g., the decay rate *c*) and functional dependencies (e.g., the activation function *G*) whose values and forms are not entirely clear or known, and complicated nonlinear terms (*Fa* and *G*(*u*)) that affect significantly the evolution of the system. The discussion below focuses on the investigation of the linear properties of the system.

## 5. The relationship between the powder-keg model 6a-6d and the WC/A class of models

We discuss below the relationship between the powder-keg model given by equations 6 and current key thermodynamic models: the class of models based on the Wilson and Cowan [1972b] formulation (the WC class), and the models based on the Amari [1977] model. These models are fundamental in the sense that, while significant efforts have been dedicated to improving the models, more recent work [e.g., Jirsa and Haken, 1996, Wright and Liley, 1995b, Jirsa and Haken, 1997, Robinson et al., 1997] is largely focused on refining the equations and may be viewed as variations of these two fundamental formulations, rather than a reexamining their foundation.

### The Wilson and Cowan [1972b] class

The thermodynamic model 6 represents a generalization of the WC class of models, similar in functionality, if not carrying exactly the same in meaning. The recipe for deriving the WC equations from system 6 is simple enough: pick a suitable form for the window *w*^*β*→*α*^, integrate in time the flux *F* (equation 6b), substitute into the kinetic energy balance equation 6a and integrate it to obtain *u*(*F*), and finally, substituting into equation 6d, obtain the firing rate as a function of the incoming energy flux.

We summarize this procedure following the choices of Wilson and Cowan [1972b], Cowan et al. [2016]. For simplicity, we assume the field comprises a single type of neurons, therefore we omit the type superscripts.

The obstacles in carrying it out reflect the differences between the two formulations. If one assumes that delays are constants and independent of axonal range, then the weighting function *w* can be factorized into spatial and temporal components

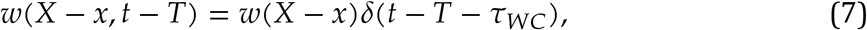

where *τ*_*WC*_is the time increment used in the discrete Wilson-Cowan equation. Equations 6a-6b become

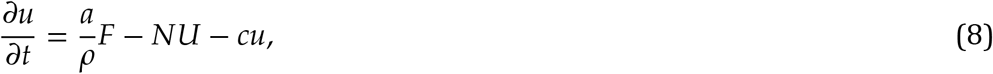

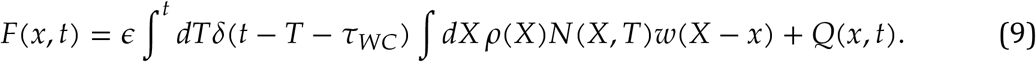

The time integration may be carried out in equation 9 to yield

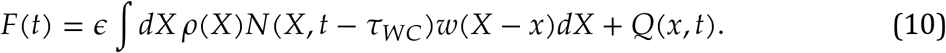

The main obstacle in this procedure becomes apparent when attempting to integrate in time equation 8. The WC formulation has no term equivalent to the *NU* term in equation 8; in general, the evolution equations for *u* and *a* obviously depend on *N* (see also equation 1) and will create a recursive algebraic dependency between *N* and *u* when substituting *u* in the equation 6d. Obviously, the evolution equations for *u* and *N* are coupled (see discussion below). We will therefore ignore the *NU* term and set *a* = 1 for now. Doing this allows for integrating the balance equation of the kinetic energy 8 to

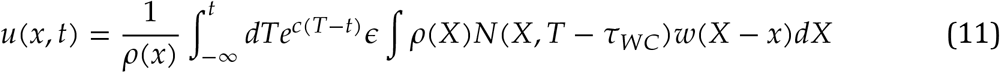

or, equivalently

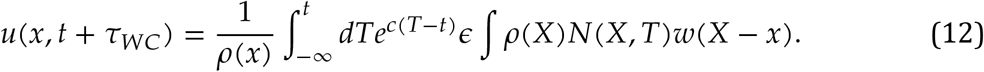

Substituting into equation 6d retrieves the functional form of the WC model, e.g., equations 7–9 in Cowan et al. [2016] (if the factors involving refractoriness and decay are ignored)

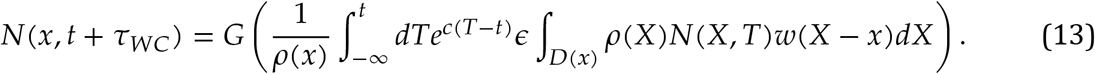

This brief derivation highlights the similarities and the differences between the model presented here and the basic Wilson-Cowen equations. Leaving aside details such as the discrete form of the latter, which of course sacrifices subgrid (cell) scales, the central difference is the description of the state of the neural field. WC models are based on the assumption that the output energy flux (firing rate) may be expressed directly as a function of the input fluxes. This assumption holds only if the flux balance does not depend on the state of the system. It is easy to see, however, that for a given fixed input flux may result in evolution trends as different as stable equilibrium (constant temperature and firing rate), catastrophic growth, or decay to zero, depending on the initial temperature of the system. One could heuristically argue that this might be the case of systems whose internal “physics” are invariant to evolution. It should be clear, however, (see figure 3) that this cannot apply to physical systems in the vicinity of a threshold-type phase transition point, and therefore to “hot” (high internal kinetic energy) neural fields, where firing depends significantly the fluctuations of the system energy. This suggests that the applicability of WC class of models is by and large limited to “cold” neural fields whose mean internal kinetic energy (temperature) is far from the firing threshold.

The absence of a state description the WC class of models may be corrected, but corrections are also limited in scope and lead to awkward behavior. For example, because the natural decay of the system toward zero temperature (term −*cu* equation 6a) cannot be introduced in a natural way, it has to be parameterized by a decay rate in the relationship between fluxes.

### Amari [1977] model

An alternative fundamental formulation that attempts to correct for the lack of a state variable is due to Amari [e.g., Amari, 1977]. The model is very similar to our equations 6, with a few significant differences. Amari introduced two new parameters, the averaged membrane potential and an excitability, and defined activation as a Heaviside function (see figure 2). The averaged membrane potential plays the role of the state variable, while excitability is assumed to be constant in time. Retrieving Amari’s model from equations 6 is straightforward. If the term *NU* is ignored and excitability parameter is constant, inserting the flux term *F* into the balance equation for *u* yields equation 1 in Amari [1977]

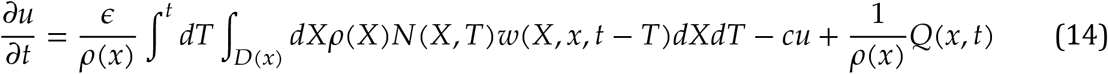

This is exactly a same functional form as given in Amari [1977], equation (1), but with a different resting state. Treating the excitability as a constant means that Amari’s model is in fact a variant of the WC class of model. This suggests that the Amari [1977] model has some (but not all) of the same limitations as the WC models. The “mean membrane potential” is not defined in the paper, and in general is hard to define when the neuron spikes. The absence of the *NU* term implies that the Amari [1977] model does not take into account the fact that spikes reset the internal kinetic energy of the neuron to zero, thus it overestimates growth and underestimates decay. A complete description of the state of the neural field requires two state variables: internal kinetic energy *u* and excitability *a*. Ignoring the time evolution of one of them (*a*) is a strong dynamical restriction. This is a drawback similar to the WC representation, albeit only partial, since *u* is used. However, the dimensionality of the phase space of the system is essentially halved.

## 6. Simplifications

The full model in Equation 6 is originally in form of integral differential equations, which is convenient for numerical simulations but poses difficulties on theoretical analysis. Under some general simplifications, we want to find a set of coupled differential equations that represent the dynamics of the original model.

For simplicity, we assume the neural field is one-dimensional and homogeneous, with negligible biological (axonal and synaptic) delays. We use the mean axonal range and the mean equivalent refractory time as units of space and time.

Then, the weighting function *w* in equation 6b is only a function of distance, *w*(*X*, *x*) = *w*(|*x*−*X*|), and substituting into equation 6b and expanding the integral formally into a Taylor series obtains

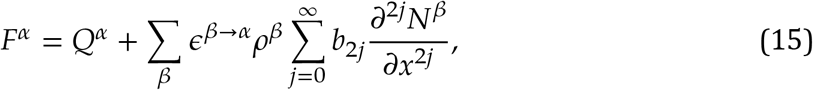

where we assume that the series is either summable, or should be interpreted as an asymptotic series, and the coefficients

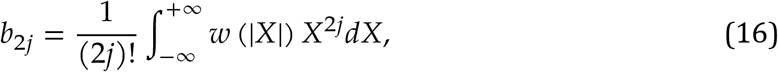

are even moments of the window *w*. In connections are uniformly distributed, i.e., the number of connections to point *x* is given by a rectangular distribution *w*(*X*, *x*) = 0.5 if |*X* ≤ *x*| 1, and zero otherwise, the constants acquire the simple form 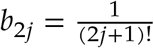. Qualitatively, higher order terms in 15 are smaller, but, as discussed in section 7, whether they are significant or not depends on the physical context.

An accurate representation of the mesoscopic refractoriness parameter is not available, but some possible simple forms are straightforward. If we assume that the mean neuron is excluded from the energy exchange process in the absolute refractory period and opens slowly post-spike, the evolution of refractoriness resemble the blue line in figure 2, bottom panel.

The standard historical convention [Cowan et al., 2016] ignores the relative refractory period and models the absolute refractory period as a rectangular distribution. The relative refractory state, however, represents a smooth transition between absolute refractoriness and full excitability: ignoring it completely is not realistic, but neither is treating it in its entirety as an absolute refractory state. It is then convenient to define the refractoriness function *r*(*t*) as an (arbitrary) decaying function with a characteristic time constant, the equivalent refractory time *τ*. Setting *t* = 0 at the beginning of the spike, the equivalent refractory time can be defined as 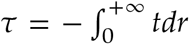 (because 0 ≤ *r* ≤ 1, the “excitability” measure 1 − *r* can be interpreted as a cumulative distribution function with mean refractory time *τ* as defined). The exact value of *τ* is somewhat arbitrary and should be determined from observational data. Throughout the discussion below we use the equivalent refractory time *τ* as the unit of time.

The Heaviside definition of refractoriness is then *r* = *H*(*τ*−*t*), where *H* is the Heaviside distribution (yellow curve in figure 2, bottom panel). Substituting into equation 6c and expanding in Taylor series on the integral over refractoriness obtains for the excitability parameter *a* the formal equation

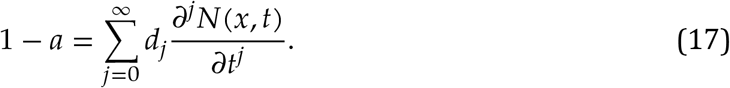

where, as above, we assume that the series symbol makes sense in some mathematical interpretation, and the integration constants *d*_*j*_ are

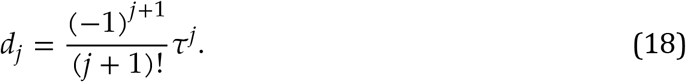

For reasons that will be discussed in detail in section 7, we propose here an alternative formulation, that models both the absolute and the relative period as an exponential decay 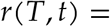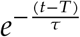(red line in figure 2, bottom panel). Substituting into equation 6c and differentiating to time obtains for *a* the equation

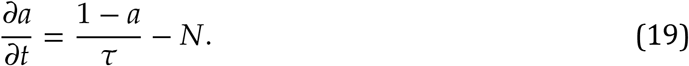

Below, we use use this form as a substitute for equation 6c.

In order to begin solving the governing equations, the functional dependency of the firing rate on the internal energy *N*(*u*), also called the “activation function”, needs to be stated explicitly. However, obtaining an physiologically accurate form of the activation function is difficult and beyond the scope of this study. The general concept of activation function dates back to Beurle and Matthews [1956] and was improved by Wilson and Cowan [1972b], who reasoned that, if all neurons in the element of area have the same mean depolarization, the firing rate is proportional the the cumulative distribution function of threshold values. Therefore, the functional form of the activation function is similar to a sigmoid. The sigmoid shape, however, is not adequate in our model for several reasons. In a randomly connected neural field the instantaneous value of the internal kinetic energy of individual neurons (blue curve in figure 2) is random (randomness of microscopic activity is a basic assumption of mesoscopic activity). While the sigmoid could be remapped to cover only the domain of our definition of the internal kinetic energy (0 ≤ *u* ≤ *U*), the goal of our model is to resolve mesoscopic time scales. The state of a neuron continuously bounces around in the interval [0, *U*], i.e., any neuron may enter refractory states and refuse to fire while accumulating the potential energy necessary for firing again. Using the sigmoid functional form in this description would imply that the neuron sub-population with zero internal energy never fires, while sub-population with *u* = *U* fires continuously.

To proceed, some assumptions need to be made (see appendix A for a discussion of the activation function). One can argue that if *u* = *U*, the firing rate is infinite; in the extreme opposite case, if *u* = 0, most (read all but a zero measure) neurons are at resting level, thus the firing rate is 0. Assuming that the activation function is monotonically increasing, a plausible functional form consistent with these constraints is

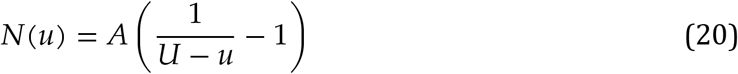

where the constant *A* is a measure of the intensity of endogenous membrane potential fluctuations (“fluctuation strength” for short). Equation 20 may be readily inverted to give

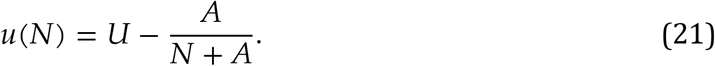

Finally, we will assume that the neural field is isotropic. This assumption implies that all odd spatial derivatives cancel, which simplifies the equations considerably, but also imposes a strong constraint that has at least two significant consequences: it enhances the diffusive character of the system, and it restricts the class of admissible solutions of equations 6, affecting in particular the wave type of solutions. Despite these drawbacks, we consider this simplification relevant for mesoscopic scales small enough to not be strongly affected, say, by boundary conditions (which are not discussed here). Nonetheless, we caution the reader that the discussion below should be regarded as relevant only for the subset of solutions satisfying this constraint, and not for the full family of solutions of the system of governing equations.

## 7. Linear analysis: single-type (excitatory) neural fields

The first step in pursuing the idea that mesoscopic collective action represents perturbative states is an investigation into equilibrium states and their stability. In this section we examine the linear properties of neural fields composed of a single neuron type (say, pyramidal cells). Below, the neural field is assumed to be under a steady, spatially uniform input, i..e., 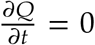 and ∇*Q* = 0. Under these conditions, the governing equations 6, written for a single-type neural field, are

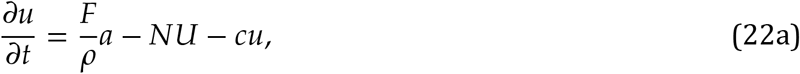

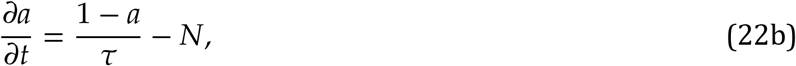

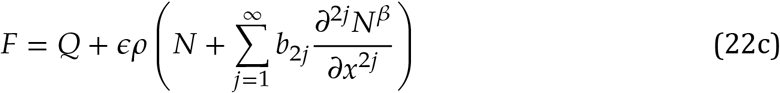

To describe perturbations around stable equilibrium states that that may vary in space, in equation 22c the energy flux was expressed the form 15. Note that, in agreements with the isotropy assumptions, only even orders of the spatial expansion are retained. In the discussion of the dispersion relation below we will prefer using for *a* equation 17, but the resulting equilibrium states are the same for both approaches.

Let *δ* ≪ 1 be a small parameter that measures the magnitude of the departure from equilibrium states, and expand the state variables in the asymptotic series

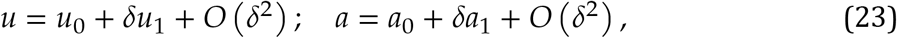

where the zero-subscripts denote the equilibrium states. For consistency, process variables *N*(*u*,*a*) and *F*(*u*, *a*) are are also expanded in asymptotic series, for example,

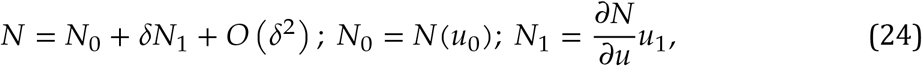

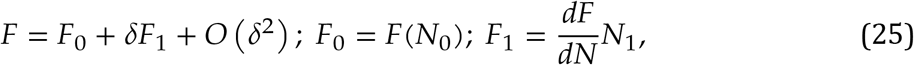

where *F* is a functional of *N*, and 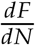 is the variational derivative.

Equilibrium states are defined here by the condition that the internal kinetic energy of the system is stationary and constant in space, 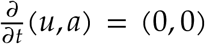 and ∇(*u*,*a*) = (0,0), therefore, the energy flux and firing rate at equilibrium are homogeneous, e.g., 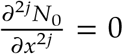. Substituting expansions into the governing equations 23–25 into the governing equations 7 and separating the powers of *δ* obtains the standard hierarchy of systems for each power of *δ*.

### 7.1. Equilibrium states

At *O* (*δ*^0^), the equations for the equilibrium state are

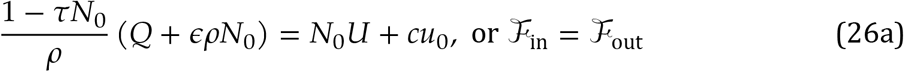

where

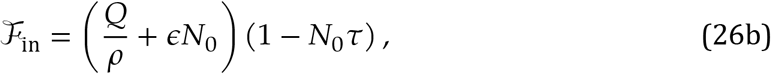

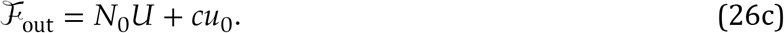

with 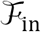 and 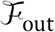 the internal kinetic energy gains and losses, respectively. Equation 26 that equilibrium states are achieved for firing rates *N*_0_such that 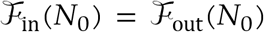. Substates stituting the pressions 26b-26c into 26a obtains the cubic algebraic equation

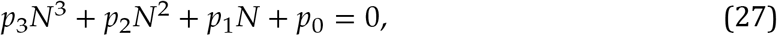

with the coefficients

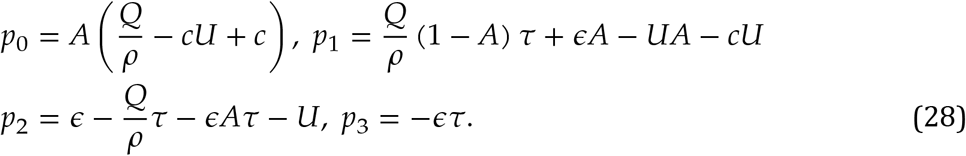

Equilibrium states correspond to the roots of equation 27. Equation 27 may have one or three real solutions corresponding to firing rates at equilibrium points, that depend on the configuration of the network. To illustrate the behavior of the system, we distinguish between two types of parameters: static parameters that characterize the physiological properties of the fields (neuron density *ρ*, decay rate *c*, threshold internal kinetic energy *U*, equivalent refractory time *τ*) and parameters that control the dynamics: connection strength (energy recaptured from a single spike) *ϵ*, and the endogenous fluctuation constant *A*. The description of equilibrium types shown in figure 4 is given for static parameters *Q*/*τ* = 0.1 and *c* = 0.5.

**Figure 4.**
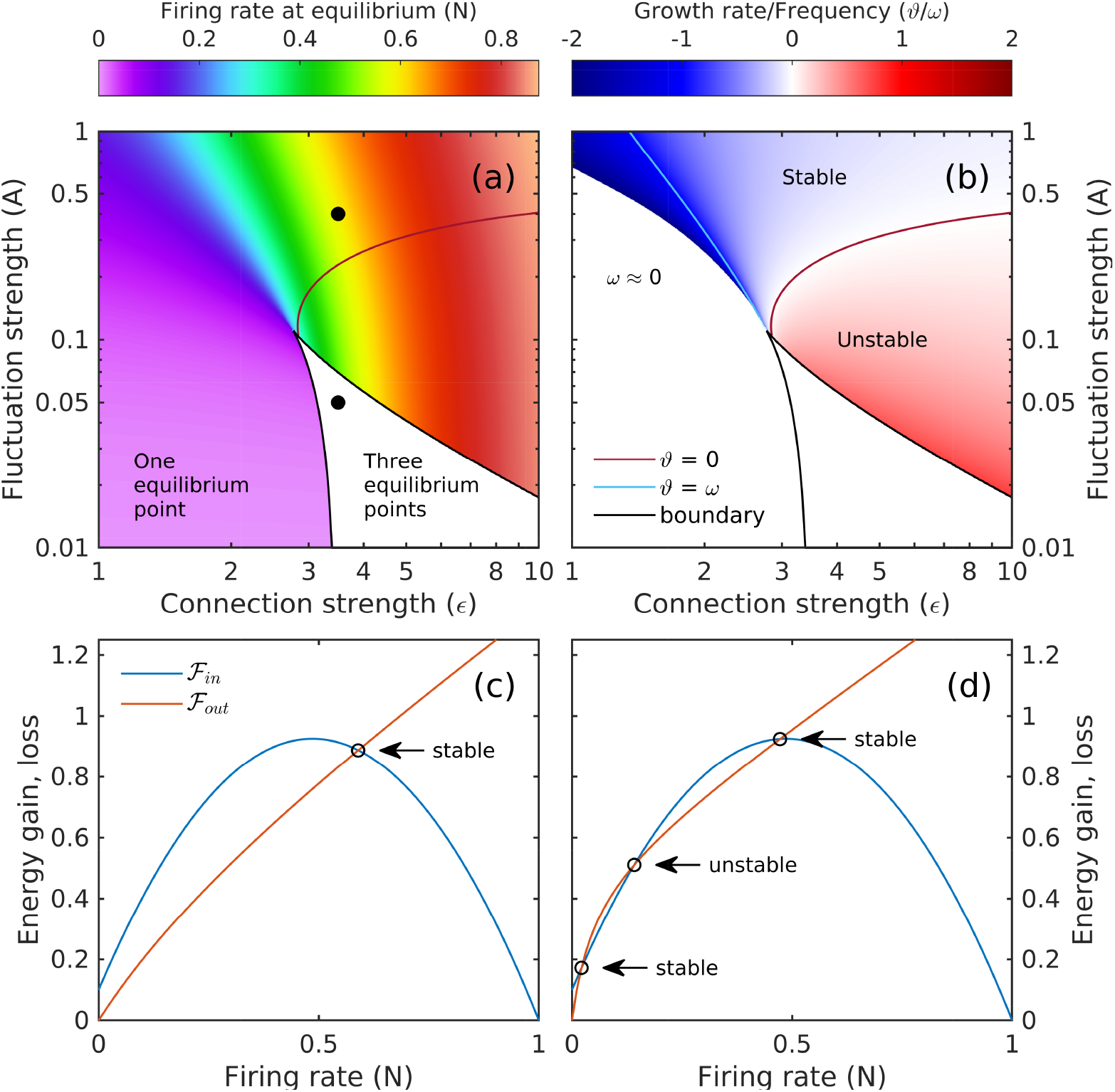
Equilibrium states under homogeneous forcing. a) Firing rate *N* at equilibrium librium (for states with one equilibrium states for cases with only 1 equilibrium state. b) The dependency of the ratio *ϑ*/*ω* at equilibrium (for states with one equilibrium point) on connection strength *ϵ* and fluctuation strengths *A*. Counterclockwise around the cusp of the domain of three equilibrium points: the frequency growth from zero to its maximum values, while the “growth” rate increases from negative values (dissipation, stable equilibrium) to positive values (true growth, unstable equilibrium). The dissipation rate equals the frequency along the blue curve, and is zero along the purple curve. c-d) Dependency of energy gains and losses (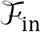 and 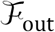) on the firing rate *N*. Equilibrium states (with firing rates *N*_0_) are realized at the intersection of the curves, i.e., 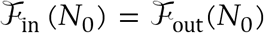. In both equilibrium cases shown (dots on panel a) parameters *Q*/*ρ* = 0.1 and *c* = 1.0, while the strength of endogenous membrane fluctuations *A* and connectivity *ϵ* are varied. c) *A* = 0.4, *ϵ* = 3.5; d) *A* = 0.05, *ϵ* = 3.5.

Single equilibrium-point configurations may correspond to different levels of firing rates *N*, as shown in figure 4.a. The dependency of the energy losses and gains (equation 26) is shown in figure 4.c. For single equilibrium points, low values of *A* and *ϵ* induce low firing rates (lower-left corner of figure 4.a), with field dynamics controlled by the external stimulation, and level of firing rate roughly proportional to the external-input level 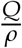. At higher values of *A* and *ϵ*, the equilibrium state is still stable, but is achieved at increasing firing rates *N*_0_ figure 4.a, (upper-right corner). As *N*_0_ increases, higher order terms in equations 26 play and increasingly important role, the relationship between *N*_0_ and external input *Q* weakens, and the stability of e equilibrium point decreases. Qualitatively speaking, as local dynamics around the fixed point gradually become unstable as *N*_0_ increases, but the nonlinearity introduced by refractory period (term *Nτ* in equation 26b) insures that the fixed points are globally stable.

Triple equilibrium states are realized for low membrane fluctuations *A* and strong connectivity *ϵ* (figure 4.a). A typical configuration of the balance of 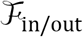 as a function of the firing rate *N* is shown in figure 4.d, with two stable points separated by a unstable one. Low values of *A* insure that low firing rates do not induce large excitability through by endogenous activity; strong connectivity insures that excitability is self-sustained at high firing rates. If stimulation or inhibition force large-enough changes in the firing rate, switching between the two stable states is possible. Because of the extreme values (low for *A*, and high for *ϵ*) we expect these cases to be rare and perhaps unrealistic, although we could not find any clear guidance in the literature about this.

Our analysis suggests that collective activity of neural populations is naturally bounded, with deviations from equilibrium state having the tendency to diffuse and average toward equilibrium. In fact, one might say that that “most” solutions are just exponential decay. Previous studies of single-type neural fields largely report only exponential decay under homogeneous perturbations, as a result, previous derivations of field equations treated the decay property as fundamental [Wilson and Cowan, 1972b, Amari, 1977]. Figure 4.b provides a qualitative representation of the extent oscillatory domain. In the (*N*, *A*) plane the oscillatory behavior is confined to relatively small domain, the white area in the neighborhood of zero-growth curve (purple). To the left (low connection strength *ϵ*) dissipation dominates (equals the frequency along the blue curve), and to the left perturbations become increasingly unstable. In a densely firing networks [e.g., Pinto et al., 2005, Trevelyan et al., 2007], refractoriness begins to play a role: if *N* is large, *a* deviates from 1 to a smaller value, which activates the nonlinear term 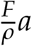 in equations 7. Because refractoriness is cumulative over time (see integral in equation 6c) it introduces in the dynamics a hysteresis effect [Cowan et al., 2016]. Due to the hysteresis, 2 a population reaches its equilibrium point form a deviated state would not just stop at the equilibrium, but the delayed effect of refractoriness changes excitability of the population so that the static equilibrium point is not dynamically stable. As a consequence, firing rate *N* is coupled to the population excitability *a* with some phase lag and the interplay between the two quantities generates a oscillatory behavior. Our model provides a mathematical formulation of this mechanism. It is worth noting that the refractory oscillatory patterns only exist in densely firing network in which refractoriness matters, that is, exist only around upper equilibrium states (figure 4.b). In comparison, dynamical patterns around lower equilibrium states only show rapid collapse because modulation from refractoriness is negligible during low firing rate.

### 7.2. Perturbations of equilibrium

At *O*(*δ*^1^), the system of equations for the leading order perturbation are

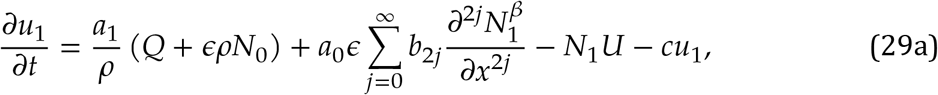

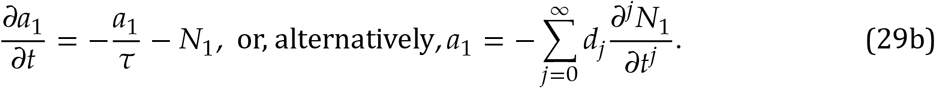

where the alternative form for *a*_1_ derives from equation 17. Equations 29 may be used to examine the stability of equilibrium states under homogeneous perturbations, or to study the dynamics of inhomogeneous perturbations (collective action).

#### 7.2.1. Homogeneous perturbations

For homogeneous perturbations 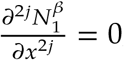, and equation 29 becomes

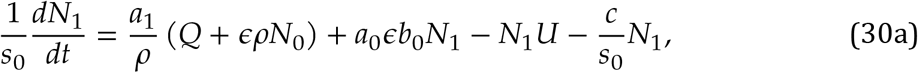

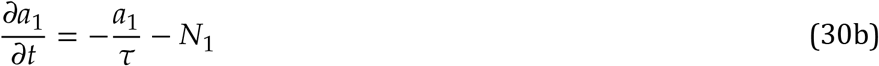

where 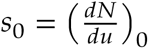. Substituting into equation 30 the standard solution *a* = *e*^*σt*^, where *σ* ∈ ℂ, with the real part *ϑ* = ℜ{*σ*} representing the growth (decay) rate, and the imaginary part *ω* = *ℑ*{*σ*} representing the frequency of oscillation, obtains

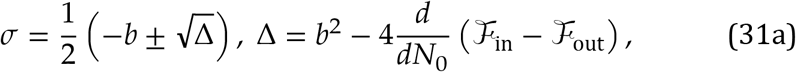

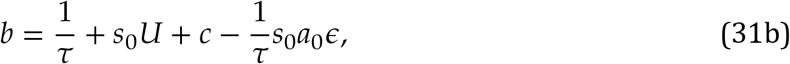

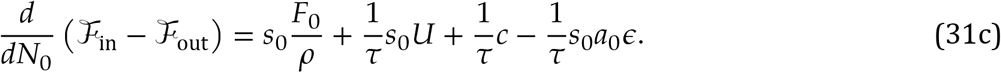

Pure growth(decay) behavior occurs if Δ ≥ 0 in equation 31a. Oscillatory perturbations may occur if Δ < 0, i.e.

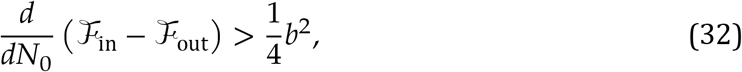

(near unstable equilibrium points - figure 4) in other words, if energy gains grow with *N*_0_ faster than losses by a margin larger than 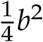. Oscillatory behavior may show growth or decay trends depending on the sign of *b* If *b* > 0, the oscillation decays as shown in figure 5.a.b for a case corresponding to figure 4.c). If the decay rate is large enough so that the inequality 32 is not possible, the dynamics is a monotonic collapse towards equilibrium (figure 5.e-f). The growth shown in figure 5.c-d corresponds to conditions near an unstable equilibrium point (figure 4.d), such that *b* < 0 and energy gains are larger than losses. As the system goes away from the equilibrium point growth rate decreases, and the trajectory of the system stabilizes along a limit cycle.

**Figure 5.**
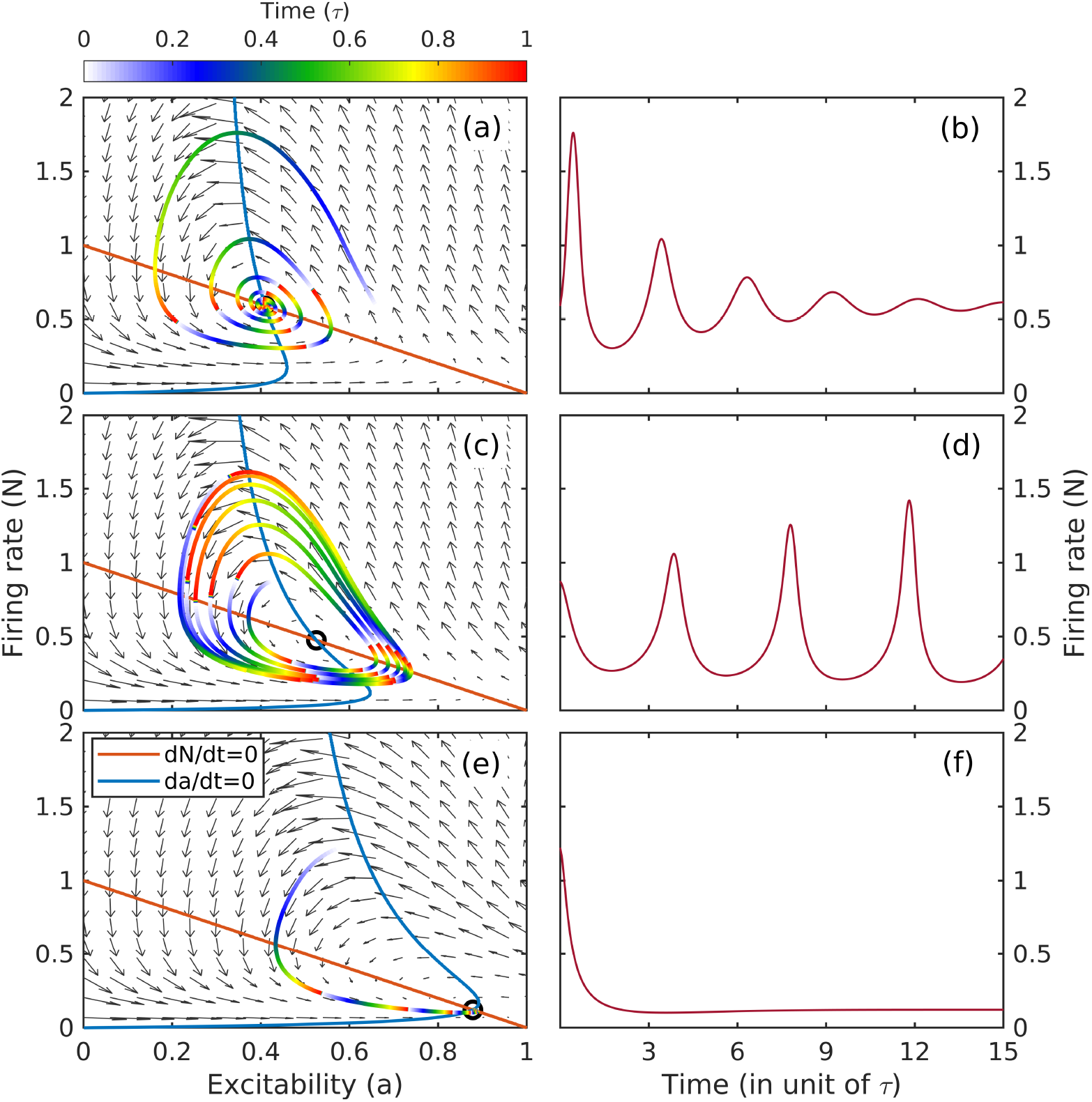
Typical oscillatory patterns of excitatory populations, *resulting from integrating the full system of equations*. Left column contains phase portraits of temporal evolution. A trace starting from an arbitrary state is shown for each case, one epoch of color map denotes one equivalent refractory period. a,c,e) Numerically integrated oscillatory patterns of firing rate. b,d,f) Time series of firing rate corresponding the right panels. a,b) Stable spiral, *b* > 0, with *ϵ* = 3.5, *Q*/*ρ* = 0.1, *c* = 0.5, *A* = 0.4; c,d) Unstable spiral, *b* > 0, with *ϵ* = 2.5, *Q*/*ρ* = 0.1, *c* = 1.0, *A* = 0.4; e,f) Stable node, *b* < 0, with *ϵ* = 4.0, *Q*/*ρ* = 0.1, *c* = 1.0, *A* = 0.4.

It is important to observe that refractoriness is the fundamental mechanism that allows for oscillatory patterns shown in the phase portraits of figure 5 arise: ignoring refractoriness is equivalent to setting *a* ≡ 1 (see equation 1) in which case equation 29 becomes a first order differential equation with no oscillatory solutions. We will therefore call these “refractory oscillations”. Refractory oscillations have periods in order *O*(*τ*), i.e., several refractory periods (e.g., figure 5), corresponding to frequency in the range of 100 Hz - 150 Hz (close to ripple frequency). When getting into spatially in-homogeneous cases, we will see spatial contribution increases slightly on the frequencies.

### 7.3. Inhomogeneous perturbations (collective action)

The analysis presented here is different if the perturbations have a non-trivial spatial structure, the spatial gradients have to be taken into account. For the sake of simplicity, it is convenient to return to Amari’s [1977] Heaviside formulation for activity (equation 17). We start, therefore, from the alternative form for the *O*(*δ*) perturbation equation 29, i.e.,

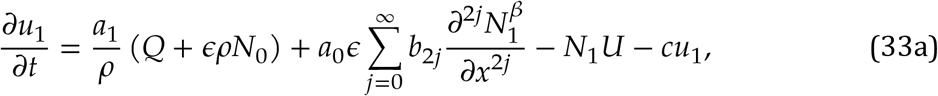

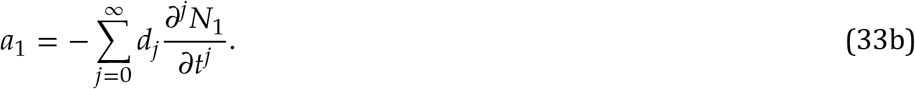

Equations 33 may be simplified to retain the internal kinetic energy *u* as the only independent variable, which obtains a single partial differential equation

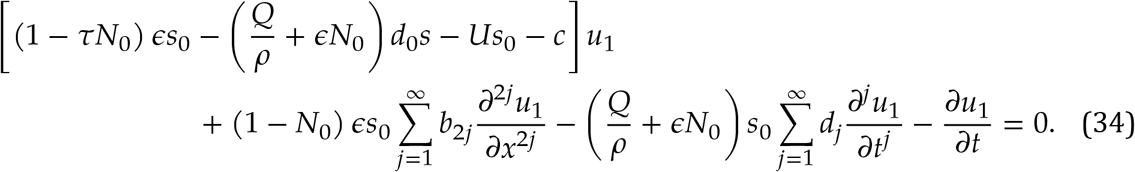

In contrast to the stability analysis in the previous section, we are interested here in identifying conditions favorable to propagating perturbations (waves). Therefore, we seek a solution in the form *u*_1_ ∝ *e*^*i*(*kx*+*σt*)^, where here *ω* = ℜ{*σ*} is the frequency and ℜ{*k*} is the wave number, and *ϑ* = ℑ{*σ*} and ℑ{*k*} are temporal and spatial growth (decay) rates. With the derivatives given by the simple rules 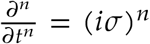 and 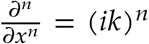 one obtains the algebraic equation

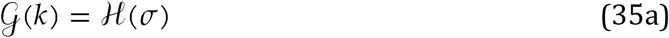

where the functions 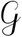 and 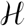are given by

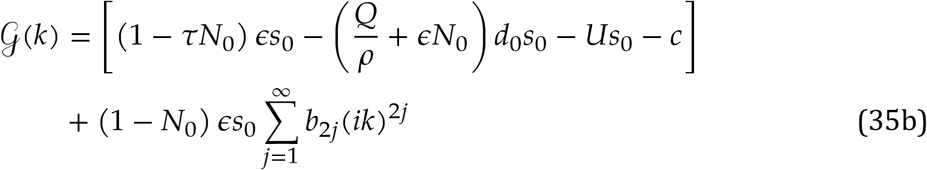

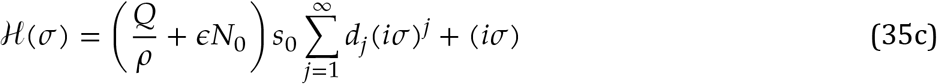

For propagating perturbations, equation 35 represents the dispersion relation [Whitham, 2011]. As a consequence of the Taylor expansions (equations 17 and 15) equation 35 contains an infinite number of terms whose significance over given temporal and spatial scales should decrease with decreasing orders of magnitude. The significance of the expansion terms for wave processes may be gauged by evaluating their contribution to the dispersion relation 35 (figure 6). While the overall trend is a monotonic decay with order in the expansion, the decay rate of terms in the temporal Taylor expansion much slower than that of the spatial terms. Keeping only the leading order approximation in equation 17, e.g., *a* ≈ (1−*τN*) is too crude to resolve wave patterns. This problem was circumvented here by introducing the exponential form of the refractoriness based on the equivalent refractory period which yielded for excitability the form in equation 19. The analysis of orders of magnitude shown in figure 6 also suggests that, because the spatial terms decay very fast, spatial coupling may be regarded as a small modulation of the temporal dynamics of homogeneous perturbations (the neural field may be approximated as a network of weakly coupled oscillators.

**Figure 6.**
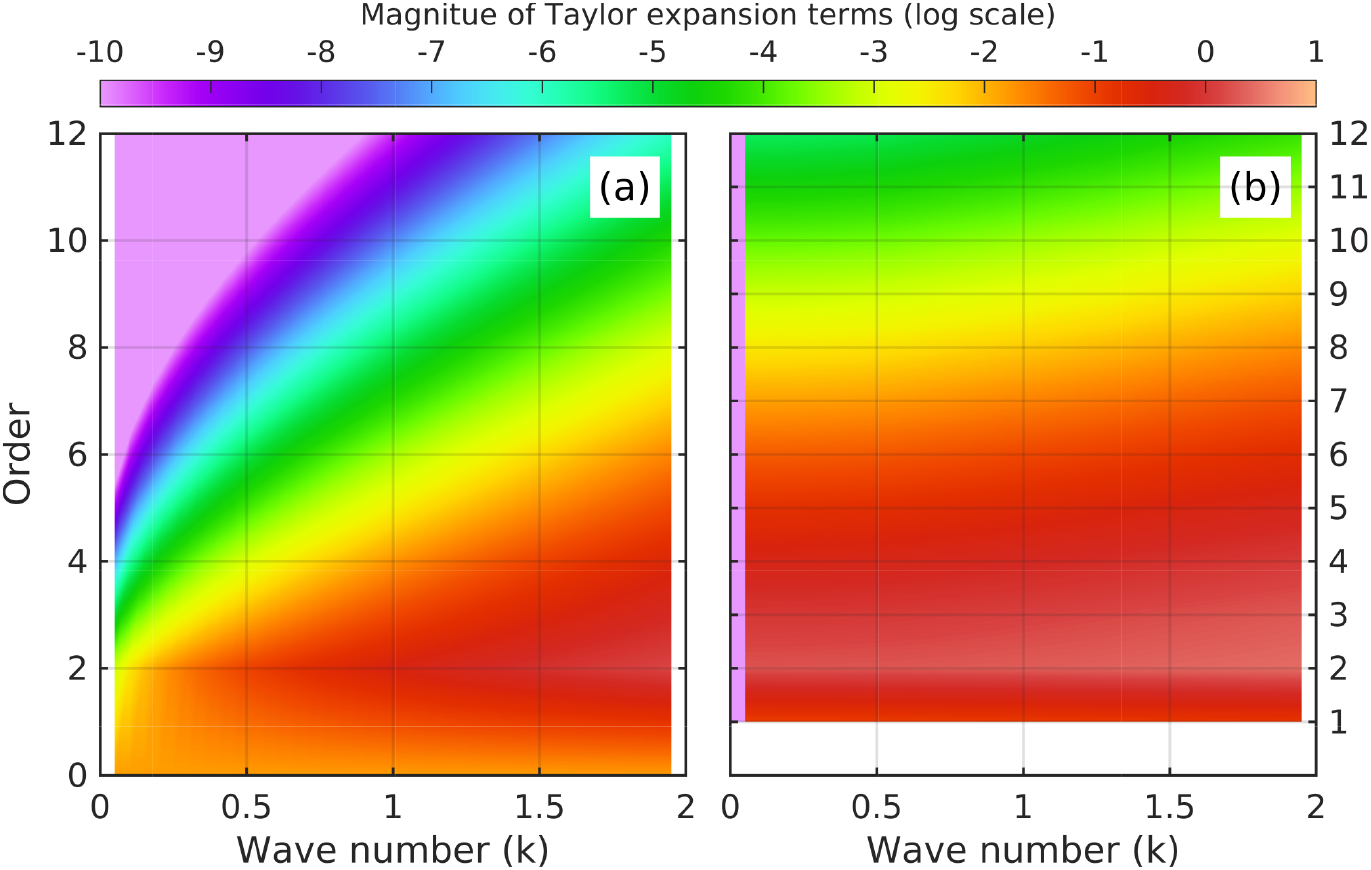
Contribution of Taylor expansion terms in the dispersion relation (equations 34 and 35): a) spatial terms (due to the isotropy assumption, only even terms are retained in the spatial expansion); b) temporal terms. This analysis provides a measure of the importance of different order approximations for the dynamics. While spatial terms decay relatively fast, the decay of temporal terms is slow and higher order terms cannot be neglected on any meaningful mesoscopic temporal scales.

To represent progressive waves, choose *k* ∈ ℝ, which implies that 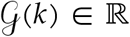 (*ik* appears at even powers), therefore 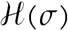 should also be real. A graphic representation of the solutions of equations 35 is shown in figure 7a-c. The resulting dispersion relation, plotted in figure 7.d, covers relatively small scales. If the wave number is 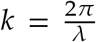, with λ the wave length in units of mean axonal range, the range plotted is between approximately 6 and 100 units. The dispersion relation *ω*(*k*) is not monotonic, but it increases overall, in a pattern similar with the decay rate, with the phase speed decreasing with *k*.

**Figure 7.**
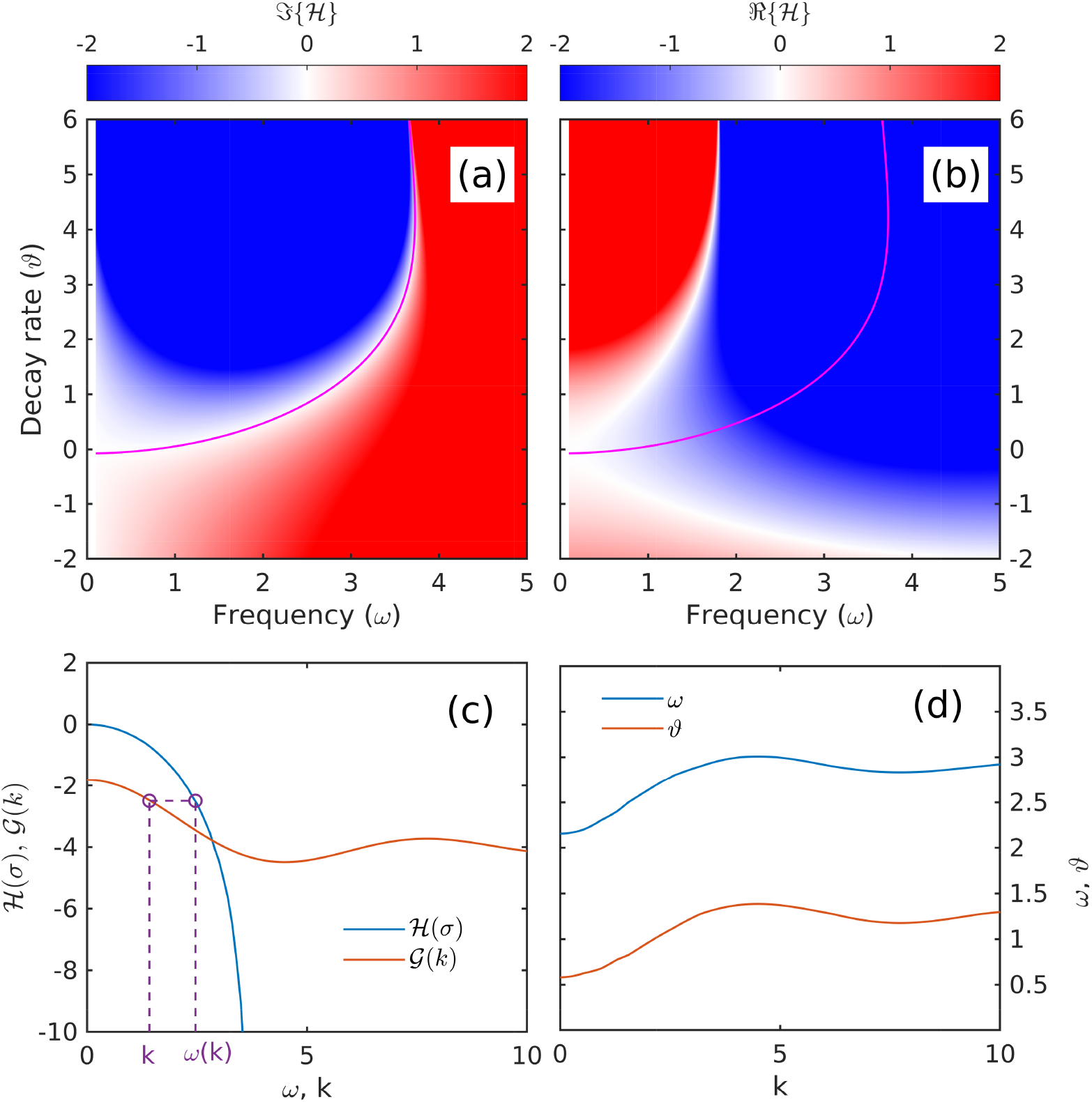
Graphic representation of the solution of equations 35 for progressive waves, for *ϵ* = 4.0, *Q*/*ρ* = 0.1, *c* = 1.0, *A* = 0.4. a) 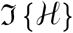, the imaginary part of 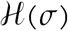 in the complex plane *σ* = *ω* + *iϑ*. The white domain (curve) in panel (a) is provides the set of all values *σ_w_* (magenta curve) such that 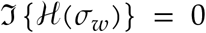. This set corresponds to waves (hence the subscript). b) 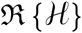, the real part of 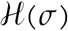 in the *σ* complex plane (the green curve is the set *σ*_*w*_). c) Graphs of 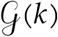 and 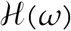, where *ω* = ℜ{*σ*_*w*_}. To find the frequency corresponding to a given value of *k*, i.e., satisfying the dispersion relation 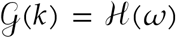, move horizontally to find 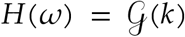, and vertically to find the corresponding *ω* (dashed purple line). d) Frequency *ω*(*k*) = ℜ{*σ*_*w*_(*k*)} and decay rate *ω* = ℑ{*σ*_*w*_(*k*)} as functions of the wave number *k*.

Due to their intimate relation with refractory oscillations discussed above for homogeneous perturbations, the wave patterns satisfying the dispersion relation 35 should be called “refractory waves”. The dynamics underlying refractory waves are similar, with propagation emerging simply as an effect of spatial coupling. The dispersion relation is monotonically increasing at low waves numbers (large wave lengths), and includes as a limiting case homo-geneous oscillations (the zero wave number has a non-zero frequency). This indicates that the lower bound of refractory-wave frequency is the frequency of refractory oscillations, which puts the frequency domain of refractory waves above the range of cortical and hippocampal ripple frequencies [Buzsáki, 2015]. The practice of detecting cortex regions with high activity by the LFP power in frequency bands associated with ripples [Ray and Maunsell, 2011] seems to support our assumption that these kind of oscillations are associated with high firing rates *N*. The behavior of the dispersion relation in the short-wave domain shown in figure 7.d also suggests that 1) the frequency band of refractory waves has an upper bound at *ω* ≈ 3; and that 2) the role of dissipation increases at small scales (the ratio of dissipation rate to frequency grows from approximately 0.25 near *k* ≈ 0 to 0.5 near *k* ≈ 4). While the plots in figure 7 are constructed for a particular set of constants, we expect them to reflect a general behavior.

The dispersion problem for excitatory networks was examined before by Meijer and Coombes [2014], who used the Wilson and Cowan [1972b, 1973] model to investigate Turing instabilities for populations with large enough refractory periods (several times of membrane time constant), looking for evidence of stationary standing or traveling solitary-wave solutions (wave “bumps”). Because the interest of their study was solitary waves, they used a numerical scheme “co-moving frame” to construct stationary solitary-wave solutions for both an equivalent delay differential model, and the original delay integro-differential model. The approach produced a dispersion-like relation between the wave speed and spatial scale, but because it refers to solitary waves, it is not a dispersion relation in the proper sense (e.g., Whitham, 2011). The proper dispersion relation was also derived by assuming a slow change of *u* over refractory period; however, the result is somewhat self-contradictory, because the solution varies on the same refractory-time scale. Because periodic waves were not the goal of the study, the dispersion relation is not discussed at length. The major contribution of Meijer and Coombes [2014] study is arguably in highlighting the essential role of refractoriness in propagating patterns of collective activity.

## 8. Linear analysis: dual-type (excitatory-inhibitory) neural fields

A dual-type neural field is of much higher interest than a single-type one, as a more realistic description of the mesoscopic dynamics of coupled excitatory-inhibitory (EI) neural fields the neocortex [Desimone and Duncan, 1995, Luck et al., 1997, Reynolds et al., 1999, Fries, 2005, Bosman et al., 2012] and hippocampus [Traub et al., 1998, Kopell et al., 2000, Bartos et al., 2007, Aton et al., 2013]. Previous studies point to inhibitory mechanisms as the main process driving rhythms in both inhibitory and excitatory-inhibitory networks. For single-type inhibitory fields the most well known mechanism is the Interneuron Network Gamma (ING; White et al., 1998, Kopell et al., 2010, Whittington et al., 2000, Wang, 2010). However, as described by [Buzsáki, 2006], a mixed population of interneurons and pyramidal cells offers complex dynamics that are capable of supporting multiple spatio-temporal patterns (for a recent review, see [Berg et al., 2019]). Recurrent connectivity between inhibitory and excitatory neurons provides the the mechanism by which a rhythmic, evolving pattern of activity can develop. The putative monosynaptic communication tends to be low latency or even synchronous [English et al., 2017, Diba et al., 2014]. Through this, it is possible to marginalize the refractory time associated with neuron to neuron communication.

Following previous studies [e.g., Amari, 1977, Jirsa and Haken, 1997, Wright and Liley, 1995b, Jirsa and Haken, 1996, Amari, 2014], we neglect for now the effects of refractory time; while we are interested in an accurate description of refractory effects, low refractoriness is adopted here as a simplification reduces the complexity of equations (population excitability becomes *a* = 1). Below, the inhibition effect is reflected by the sign of the energy recaptured by the field from firings by inhibitory neurons: we assume *ϵ*^*E*→*E*^ > and *ϵ*^*E*→*I*^ > 0, but *ϵ*^*I*→*I*^ < 0 and *ϵ*^*I*→*E*^ < 0.

The linear analysis of the equations for dual-type neural fields follows the same steps as used in Section 7. Starting from the governing equations 6, we apply the simplifications introduced in section 6, and neglecting the refractory terms, the governing equation may be written as

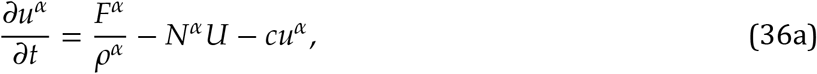

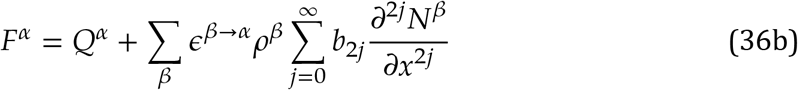

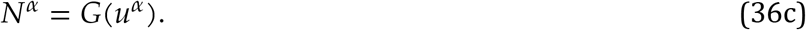

where *α* = *E*, *I* for excitatory and inhibitory neurons, respectively. The governing equations were simplified further by assuming that parameters *c*, *b*_2*j*_, and *U* do not depend on neuron type. Expanding as before the variables in the asymptotic series

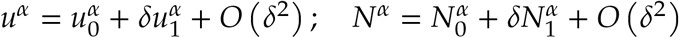

and substituting into the governing equations produces the two systems of equations for the equilibrium states and for the leading order perturbations.

### 8.1. Equilibrium states

At *O*(*δ*^0^), the equations for the equilibrium state are:

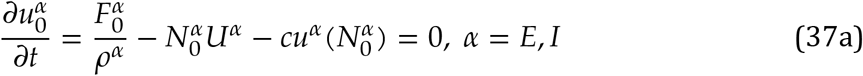

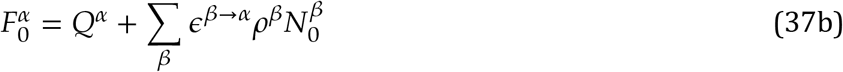

Taking the firing rates *N^α^* as free parameters, the solutions of equations 37a may be obtained graphically by examining the intersections surfaces ∂*u^α^*/∂*t* as functions of *N^α^* with the zero plane (figure 8, left panels). A visualization of the equilibrium states as the intersection of the two curves obtained this way is shown in figure 8. As before, coupled dynamics of excitatory and inhibitory neural fields problem depends on parameters *c,A,ϵ,ρ* and *Q* but the number of parameters doubles. Because an exhaustive exploration of the parameter space is beyond the scope of this discussion, we assume again that the important dynamical parameters are *A* and *ϵ*^*I*→*E*^ or *ϵ*^*E*→*I*^. As suggested by the analysis of a single-type neural field, *A* plays an important role in equilibrium bifurcation, and parameters *ϵ*^*I*→*E*^ or *ϵ*^*E*→*I*^ (connection strengths between inhibitory and excitatory neurons) describe the effect of inhibition, which is the interesting point in dual types of neurons: if either *ϵ*^*I*→*E*^ or *ϵ*^*E*→*I*^ cancel, the field defaults to the single-type neural field, discussed in section 7. The rest of the parameters are assumed kept constant at the (arbitrary) values *ϵ*^*E*→*E*^ = 4, *ϵ*^*I*→*I*^ = 0, *Q*^*E*^/*ρ*^*E*^ = 0.1, *Q*^*I*^/*ρ*^*I*^ = 0 *c* = 0.5, *A* = 0.4.

**Figure 8.**
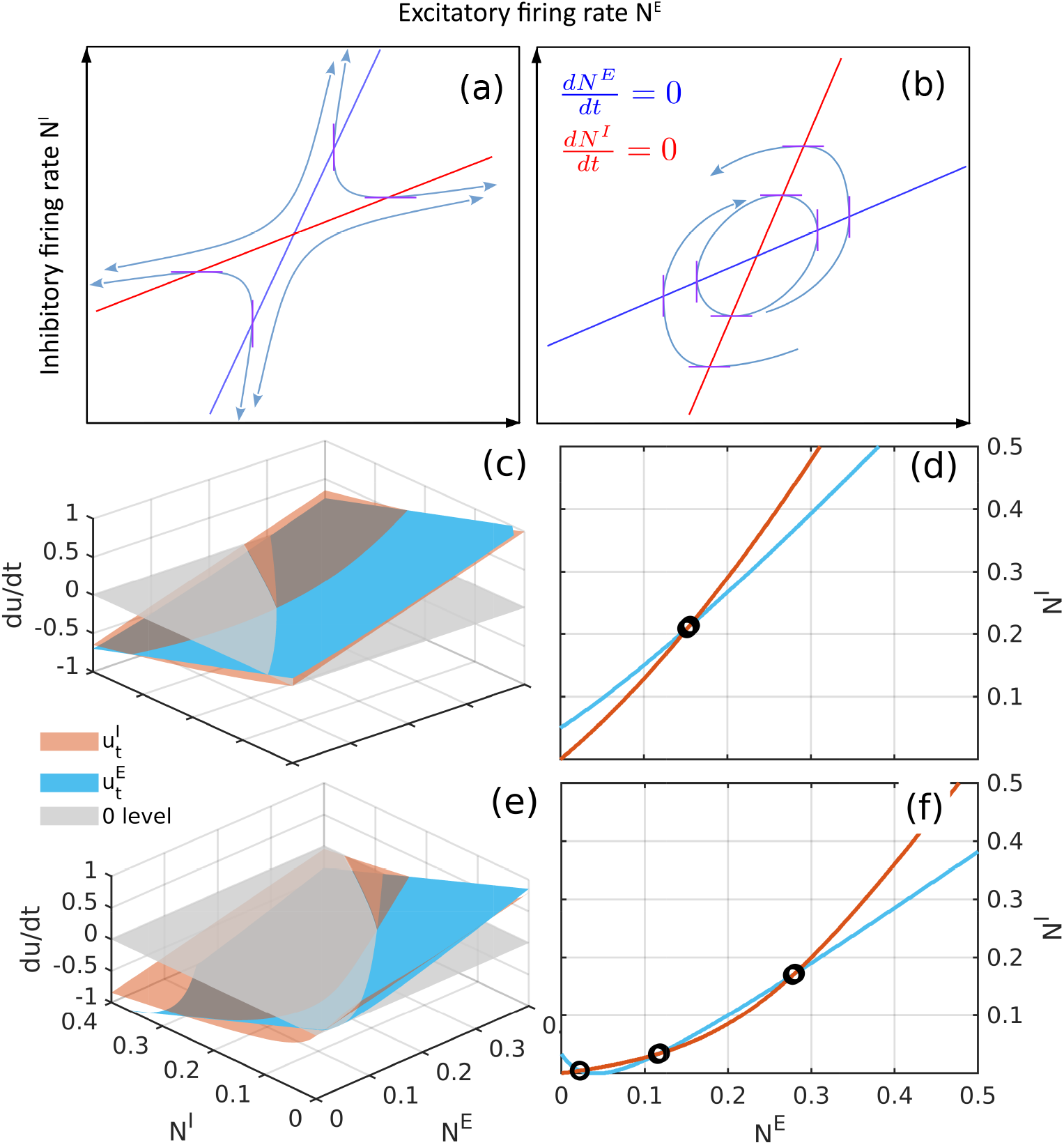
Typical equilibrium states of coupled excitatory and inhibitory populations. a-b) Possible configurations of curves 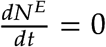 (blue), and 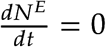 (red). The angle between the direction of the associated the vector field and the *N^E^* axis is 90 degrees along the blue curve and 0 degrees along the red curve, and ≠ 0,90 degrees everywhere else. c,e) An illustration of equilibrium points as intersections of the surfaces 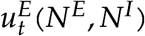 (blue) and 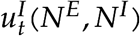 (red) and the zero surface (gray). d,f) Illustration of the equilibrium points as intersections of the curves 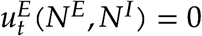 and 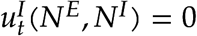 (curves representing the intersection of the blue and red surfaces with the gray one). Panels (c.d) correspond to a single equilibrium fixed ponit (*ϵ*^*E*→*E*^ = 4, *ϵ*^*E*→*I*^ = 2.4, *ϵ*^*I*→*E*^ −2.5, *ϵ*^*I*→*I*^ = 0, *Q^E^*/*ρ^E^* = 0.1, *Q^I^*/*ρ^I^* = 0 *c^E^* = 0.5, *A*^*E*^ = *A^I^* = 0.4). Panels (e.f) correspond to a case with three equilibrium points (*ϵ*^*E*→*E*^ = 4, *ϵ*^*E*→*I*^ = 2, *ϵ*^*I*→*E*^ = −3, *ϵ*^*I*→*I*^ = 0, *Q^E^*/*ρ^E^* = 0.1, *Q^I^*/*ρ^I^* = 0 *c^E^* = *c^I^* = 0.5, *A^E^* = *A^I^* = 0.05)

### 8.2. Perturbations of equilibrium

At *O*(*δ*^1^), replacing 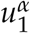 in leading order by 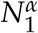, the perturbation the equations are,

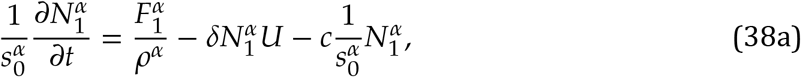

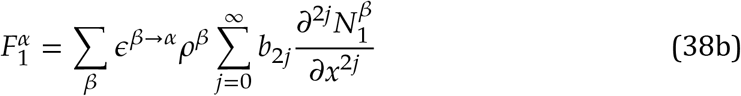

where 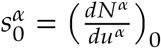, and 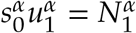 if *δ*^2^ terms are ignored. For homogeneous perturbations, equation 38 reduces to

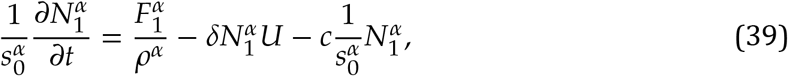

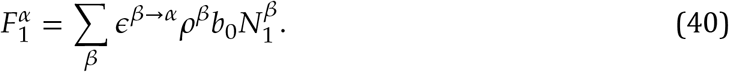

#### 8.2.1. Homogeneous perturbations

As before, for stability analysis, solutions are sought in the form *N*^*α*^ = *C^α^ e^σt^*, where *σ* ∈ ℂ, with the real part *ϑ* = ℜ{*σ*} representing the growth(decay) rate, and the imaginary part *ω* = ℑ{*σ*} representing the frequency of oscillation. We assume that the two neuron populations have the same type of dynamics (growth rate, frequency), but allow for different amplitudes and phases, represented by *C^α^* ∈ ℂ. Therefore, the phase lag between the two populations is defined as

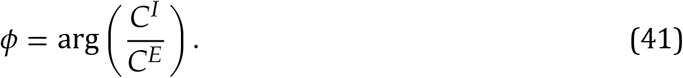

With these notations, straightforward algebra (see details in appendix, section B) obtains

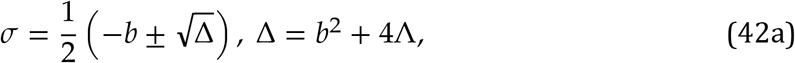

where

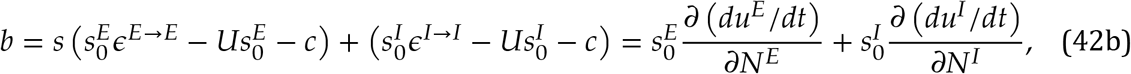

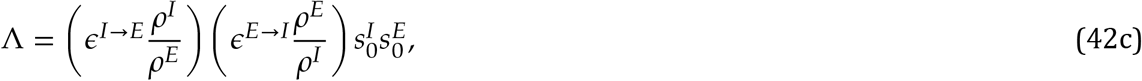

and the phase lag is

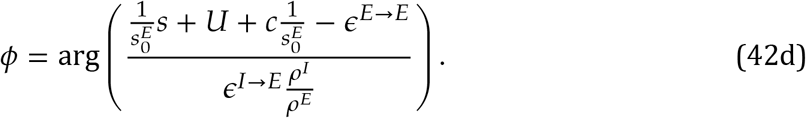

As before, oscillatory patterns correspond to Δ < 0 in equation 42a. In contrast with the case single-type field, for EI fields the term 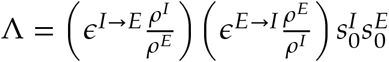 always negative because of the inhibitory effect (*ϵ*^*I*→*E*^ < 0), therefore oscillations are naturally available. The relation 42b between *b* and the partial derivatives of the rate of change of the internal kinetic energy provides a useful tool to understand the stability of the equilibrium states. From figure 8, the slope 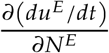 of the blue surface along the *N^E^* axis) can be either positive or negative, while the slope 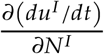 of *du*^*I*^/*dt* in the *N*^*I*^ direction is naturally negative (because 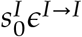 is always negative thus 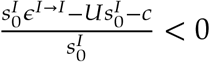 as well). Therefore, *b* can be either either positive or negative, implying that growth and decay are both mechanistically supported around the equilibrium states. The phase portraits sketched in figure 8.a-b, have the geometric constraints that the vector field has to be vertical along the curve 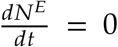(blue), and nowhere else, and and horizontal along the curve 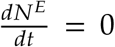(red), and nowhere else. However, because the actual direction of the flow is not specified, if the slope of the blue curve is larger than the slope of the red curve, the equilibrium point is an unstable saddle point; if slope of the blue curve is less than slope of red curve (figure 8.b) the equilibrium point can be either a center, or a stable spiral, or an unstable spiral.

Figure 9 includes several visualizations of dynamical patterns in the phase portraits. In the neighborhood of stable equilibrium states, if Δ < 0 and the connectivity *ϵ*^*E*→*E*^ is weak enough, the interaction between excitatory neurons may not be enough to maintain oscillatory amplitudes (*b* is not likely to be a positive value), and a decaying oscillatory pattern arises (figure 9.a.b). Specially when the decay rate *b* is small enough, almost no oscillatory pattern will be seen and the dynamics is a nearly monotonic collapse towards equilibrium (figure 9.e-f). Near unstable equilibrium states (figure 9.c-d), if Δ < 0 and *b* > 0, the interaction between excitatory and inhibitory neurons amplifies oscillatory amplitudes.

**Figure 9.**
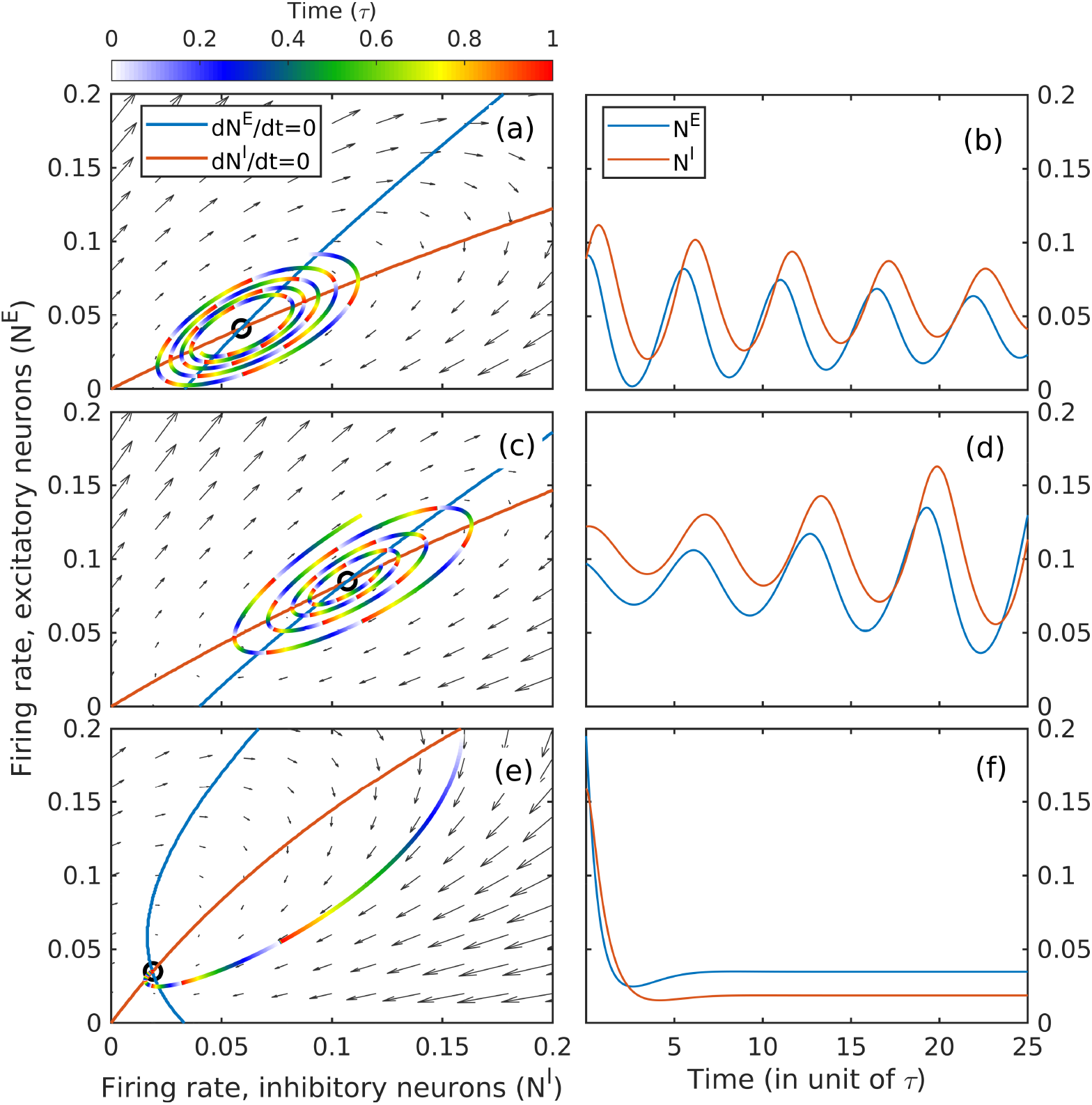
Typical oscillatory patterns of coupled excitatory and inhibitory populations. Left column contains phase portraits of temporal evolution. A trace starting from an arbitrary state is shown for each case, one epoch of color map denotes one equivalent refractory period. Right column contains numerically integrated oscillatory patterns of firing rate for both cases with *b* < 0, parameters are and respectively *ϵ*^*E*→*E*^ = 4, *ϵ*^*E*→*I*^ = 3, *ϵ*^*I*→*E*^ = −3, *ϵ*^*I*→*I*^ = 0, *Q*^*E*^/*ρ^E^* = 0.1, *Q*^*I*^/*ρ^I^* = 0 *c^E^* = *c^I^* = 0.5, *A^E^* = *A^I^* = 0.4 and *ϵ*^*E*→*E*^ = 4, *ϵ*^*E*→*I*^ = 3, *ϵ*^*I*→*E*^ = −3, *ϵ*^*I*→*I*^ = 0, *Q*^*E*^/*ρ^E^* = 0.1, *Q*^*I*^/*ρ^I^* = 0 *c^E^* = *c^I^* = 0.5, *A^E^* = *A^I^* = 0.4

In contrast to single-type excitatory neural fields, which support refractory oscillations only at high firing rates, oscillatory patterns exist in EI fields even when firing rate is low. This indicates that the generating mechanism of the homogeneous oscillations of EI fields shown in figure 9 relies on interaction between the two types of neurons. These oscillations will be referred to as “interactive oscillations”. While refractory oscillations may exist only in densely firing networks (high *ϵ*), EI-type of interactive oscillations may be generated at low firing rates, i.e., near lower-activity equilibrium states. Qualitatively, increased activity of excitatory neurons increases the internal kinetic energy of connected inhibitory population. Cumulative hysteresis effects on the inhibitory internal kinetic energy triggers delayed activation rates of inhibitory neurons, which, in turn inhibit excitatory activity. The firing rate of excitatory population drops below equilibrium, but, as the kinetic energy of the inhibitory population also drops below equilibrium, the excitatory population recovers the ability of high firing rates. Our model provides a mathematical description of this mechanism, in contrast with other models, that rely on structural delays to generate waves (e.g., axonal delays in Jirsa and Haken, 1996, 1997 and Wright and Liley, 1995b; update delay in Cowan et al., 2016). In fact, our simplified model is able to treat delays as negligible and still resolve oscillations.

The examples shown in figure 9 suggest that the time scales (periods) of interactive oscillatory patterns (decided by the discriminant in equation 42a), are similar in magnitude to refractory oscillations (several refractory periods, i.e., frequencies between 80 Hz to 130 Hz). When we get into spatially in-homogeneous cases, we will see spatial contribution decreases slightly on the frequencies. The observation that interactive oscillations may be generated at lower firing rate is consistent with in-vivo observations of gamma waves [Ray and Maunsell, 2011]. This suggests that the EI interactive oscillations generated by coupled excitatory and inhibitory fields in sparsely firing networks might provide a mathematical basis for understanding the fundamental oscillatory frequency identified as gamma.

### 8.3. Inhomogeneous perturbations (collective action)

As before, following the “progressive wave” convention, we look for solutions in the form *N^α^* = *C^α^ e*^*i*(*kx*+*σt*)^, *α* = *E*, *I*, where real values represent oscillations and imaginary values represent decay of growth. We also assume that the mesoscopic activity of both excitatory and inhibitory populations is characterized by the same spatial and temporal structure. Substituting into equation 38 obtains the algebraic equation (dispersion relation for waves; see details of the algebra in appendix C)

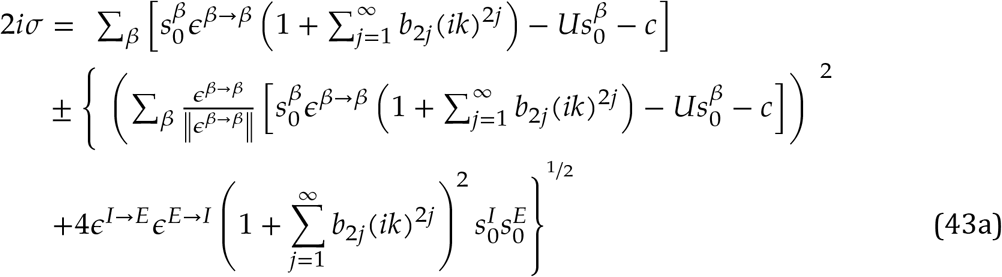

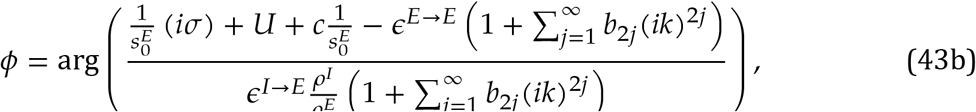

Similar as our interest in temporal dynamics of excitatory neurons, dynamical patterns of temporal interactions between excitatory and inhibitory neurons are also studied. The wave frequency *ω* and growth rate *α* as a function of real wave number *k* are plotted in Figure 10 for an illustrative case.

**Figure 10.**
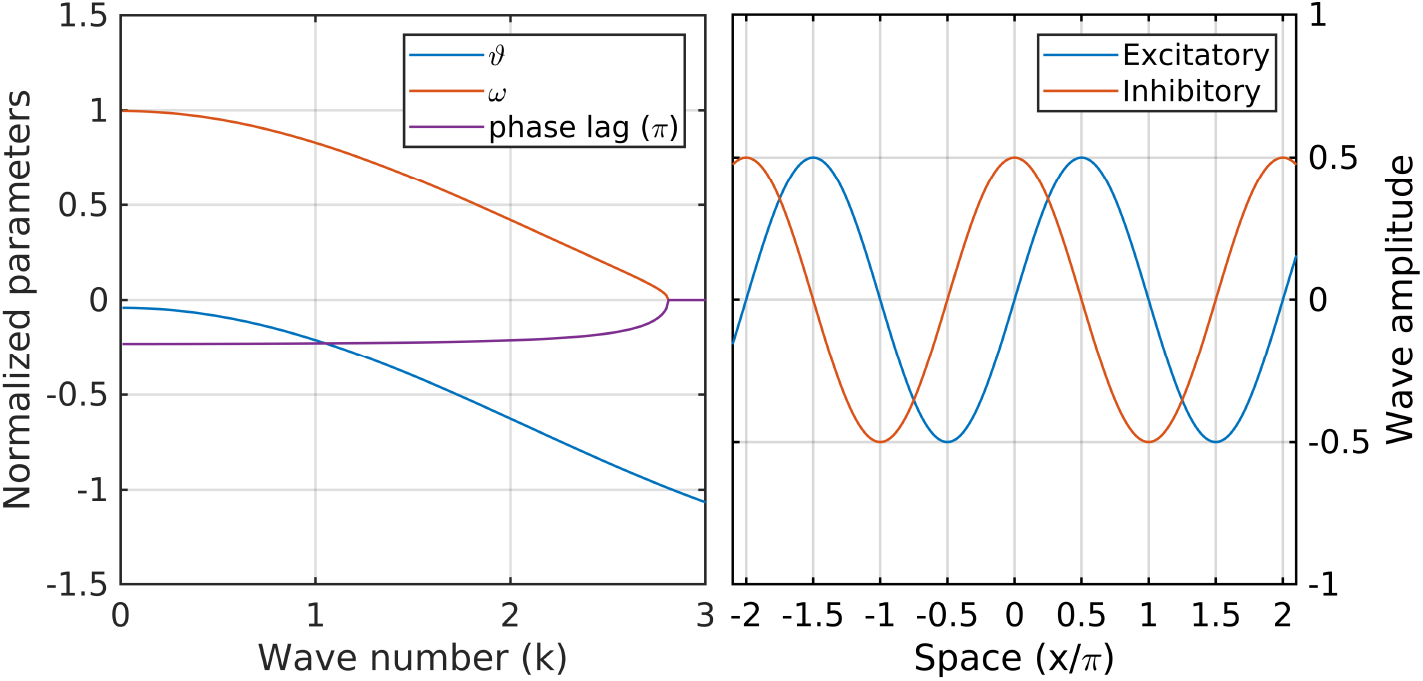
Left panel: A typical dispersion relation and phase lag of coupled excitatory & inhibitory neurons. Parameters for this case are: *ϵ*^*E*→*E*^ = 4, *ϵ*^*E*→*I*^ = 3, *ϵ*^*I*→*E*^ = −3, *ϵ*^*I*→*I*^ = 0, *Q*^*E*^/*ρ^E^* = 0.1, *Q*^*I*^/*ρ^I^* = 0 *c^E^* = *c^I^* = 0.5, *A^E^* = *A^I^* = 0.4; Right panel: Schematic wave form of linear interactive waves.

This sort of waves shown in the dispersion relation are called interactive waves analogous to interactive oscillations in the homogeneous case. Interactive waves (figure 10.a) show crests of inhibitory activity lagging behind excitatory activity (in the case shown, the phase lag is approximately *π*/4). Qualitatively, the wave pattern may be described as a hysteresis-driven alternation of highs and lows of excitatory activity, which triggers a delayed increase of internal kinetic energy in locally connected inhibitory population. Thus, lagging inhibitory activity suppresses local excitatory population, and the cycle repeats itself.

The parameters characterizing the dispersion relation of interactive waves are shown in figure 10. Remarkably, the frequency is monotonically decreasing with increasing amplitudes, but the wave character of these patterns also depends on the dissipation rate, which increases with the wave number. The domain of interactive waves is effectively cut off in the neighborhood of *k* ≈ 1; above this value, the dissipation rate becomes comparable, implying that the perturbations decay too fast to qualify as oscillations in time. Following the same reasoning as for single-type neural fields, this suggests that interactive oscillations (which could be identified as zero wave-number interactive waves) provide an upper bound for the frequency range of interactive waves, consistent with gamma frequencies in sparsely-firing networks.

## 9. Discussion

Our prior interpretation of spectra and bispectra of hippocampal LFP suggests that mesoscopic collective activity is a perturbation of an background (equilibrium) state that displays the fundamental features of a turbulent system: weak nonlinearity [Sheremet et al., 2019b], stochastic behavior [Freeman, 2000b,a, Sheremet et al., 2018b, Zhou et al., 2019], and weak dissipation. To investigate further this hypothesis requires theoretical and numerical models capable of describing activity of a large populations of neurons. Although mesoscopic activity has been the focus of considerable research, the key thermodynamic models, due to Wilson and Cowan [1972b, 1973], Cowan et al. [2016] and Amari [1977], have drawbacks significant enough to consider revisiting the formulation of the governing equations.

Here, we present the derivation of a thermodynamic model for mesoscale collective action based on the fundamental assumption that the mean neuron in a neural field is characterized by two different stages of evolution: 1) a sub-threshold stage, in which the neuron is at “microscopic” equilibrium, well described by the potential averaged over the surface of the cell membrane; and 2) a transitional stage, corresponding to the potential spiking, in which a large electric pulse propagates along neural membrane. The latter stage has the remarkable property that the neuron is for a short period of time essentially unresponsive to stimuli (absolute refractory state). From a thermodynamic perspective, the former stage is characterized by an internal kinetic energy which could be defined as proportional to the averaged membrane potential. However, the averaged membrane potential is not well defined during the firing process, and the state of the neuron is ill defined. This suggests distinguishing between two types of energy: a potential energy, released during a spike, and the internal, sub-threshold kinetic energy, that serves as the trigger for a spike. From a thermodynamics perspective, internal kinetic energy is a state variable, i.e., characterizing the state of the neuron. In contrast, the energy captured from the potential energy released by a firing is a process variable, e.g., similar to heat fluxes in classical thermodynamics.

The thermodynamic formulation based on these considerations on the dynamics of the “leaky integrate-and-fire” neuron model is essentially the powder-keg paradigm. The “temperature” of a keg plays the role of internal kinetic energy: if it exceeds a threshold, it triggers the explosion of the keg, i.e., the release of the potential energy. Some of the energy released is recaptured by the system, increasing locally the temperature, as well as providing temporal (oscillatory) organization. From a thermodynamic perspective, a large collection of powder kegs is described by two state variables: the excitability and the internal kinetic energy of the element of volume of the neural field. The process of neurons firing is treated as a process variable involved in the energy exchange of the system with its environment. The formalization of this concept leads to a system of integro-differential equations that may be seen as a generalization of the Wilson and Cowan [1972b, 1973] and Amari [1977] models, with the main advantage being the explicit evolution equations for the two state variables.

We examined linear approximations of the governing equations for single-type (excitatory) and dual-type (excitatory-inhibitory) neural fields. Both cases exhibit states with internal kinetic energy balance that translate into single- or triple-point equilibrium states. Our analysis agrees with previous observations (e.g., Meijer and Coombes, 2014, Coombes et al., 2014, Muller et al., 2018a) that the refractoriness property of the system, i.e., the existence at any time of a fraction of neural population that is “disabled” and cannot fire, is a crucial element in the generation of oscillatory behavior. In single-type neural systems, this ability is provided by the natural refractory state of a neuron, with the direct consequence that temporal scale of both homogeneous and inhomogeneous oscillations is of the order of the refractory period. We call these “refractory oscillations/waves”. In dual-type systems, the inhibitory component can take over this function and the system can support oscillations even if the refractory period of individual neurons is ignored. We call these “interactive oscillations/waves”. This property is at the root of the major difference in the linear behavior of the two types of systems. The dynamics of single-type excitatory neural fields are naturally decaying, with all equilibrium states globally stable, and with oscillations occupying a “small” domain in the phase space (figure 4.b), typically corresponding to high firing rate. In contrast, dual-type (excitatory-inhibitory) neural fields support oscillations at much lower activity levels, and are intrinsically more unstable, with globally unstable states possible.

In interpreting the results of the linear analysis it is important to note that 1) the discussion refers to the linear analysis of a simplified version of the governing equations 6 and not of the full equations (this is particularly relevant for wave solutions, given the strong isotropy constraint imposed); and 2) that, although the analysis of the linear system is essential for understanding the nonlinear behavior of the system, it does not provide an interpretation of the spectral shapes observed (any spectral shape may correspond to a solution of the linear system). Nonetheless, isotropic results should be relevant at least for small enough mesoscopic scales (e.g., gamma oscillations and ripples); and linear considerations do provide an interpretation of the local dynamics at different scales.

Withe these reservations, and assuming that the model presented here has any relevance for the interpretation of LFP measurements, several suggestions seem to emerge:

1. The linear analysis shown provides a representation of processes that occupy the ripple and gamma frequency bands. Singe-type neural fields support refractory oscillations and waves only at high firing rates (*N*), consistent with observations of “replay during ripples” [Kudrimoti et al., 1999].
2. The theta rhythm does not satisfy the dispersion relation 10 (dissipation of interactive waves is too strong at theta scale), implying that theta cannot propagate as a free wave in the hippocampus, hence it has to be an externally forced oscillation. While this is consistent with the global nature of theta, observations [e.g., Lubenov and Siapas, 2009] do show that theta has a well defined direction of propagation in the hippocampus, and therefore does not satisfy our isotropy constraint. It is therefore possible that theta simply does not belong to the family of isotropic solutions discussed here. Either way, the analysis presented here suggests that global theta forcing may play a major role in modulating key parameters of the system: internal kinetic energy and excitability (refractoriness) levels, and thus in maintaining equilibrium states, and providing the increased activity necessary to sustain mesoscopic collective action.
3. Previous nonlinear analysis [Sheremet et al., 2019b] suggests that gamma oscillations reside preferentially in the theta trough (e.g., theta-gamma biphase ≈ 180 degrees). This is consistent with the “linear” analysis: the trough of theta corresponds to locally higher forced activity levels (higher external input *Q* in our model). In the linear model, increased energy input decreases the stability of the equilibrium state, facilitating mesoscopic oscillations.
4. Revisiting the schematic spectra in figure 1, our analysis suggests that the gamma frequency band is occupied by interactive processes, possibly waves, bounded above by nearly-homogeneous oscillations (see the dispersion relation in figure 10). In the upper frequency bands, probably dominated by refractory processes, the role of waves and oscillations reverts, with oscillations having lower frequencies than waves. If theta is considered strictly as a forcing term, the increase of gamma power with theta is consistent with the increase of the oscillation amplitude with the forcing.

The model presented here comes with the overall implicit - and parsimonious-assumption that brain activity may be described within the framework of thermodynamics, providing a background to understand the physics by which the brain organizes behavior. The ubiquity across species and brain regions of isotropic and homogeneous mesoscale neuronal structures [Lorente de No, 1938, Parent and Hazrati, 1995, Marder and Bucher, 2001, Garamszegi and Eens, 2004, Apps and Garwicz, 2005, Mante et al., 2013] suggests the existence of a “universal computational principle”. Freeman and Vitiello [2010] hypothesize (citing Lashley, 1942), that mesoscale processes are the essential cognition step of abstraction and generalization of a particular stimulus to a category of equivalent inputs, “because they require the formation of nonlocal, very large-scale statistical ensembles (our emphasis)”. As often argued [e.g., Freeman, 2000a, Frisch, 2014, Edelman and Gally, 2001], physical processes underlying cognition are expected to resemble biological processes, with no design and no a priori function [Edelman and Gally, 2001]. Frisch [2014] notes that “biological systems have an intrinsic ability to maintain functions in the course of structural changes”, such that “specific functions can obviously be constituted on the basis of structurally different elements, a biological property that is referred to under the term degeneracy [Edelman and Gally, 2001]”. It is possible that mesoscopic collective action is the basis of the “universal computational principle”. As computational support, mesoscopic collective action has significant reconfiguration potential, especially under a priori unknown conditions [Sussillo and Abbott, 2009]. Understanding mesoscopic activity dynamics may be the first step toward understanding the elusive process of brain integration.

## Acknowledgments

This work was supported by the McKnight Brain Research Foundation, and NIH grants. Grant Sponsor: National Institute on Aging; Grant number: AG055544 and Grant Sponsor: National Institute of Mental Health; Grant Number: MH109548.

## Appendix A. The activation function and the positive-definite character of the *u* and *a*

Here, we discuss the hypotheses and approximations used in the derivation of the activation function used in this study. Although the powder-keg model belongs to the Wilson and Cowan [1972b, 1973] and Amari [1977] class of models, the definition of essential variables such as the firing rate *N* and the state variables *u* (mean internal kinetic energy) and *a* (excitability) is different enough to require a re-examination of the activation function. Because the focus of this study is to construct the model and examine its basic properties, much of the derivation presented below is driven by the need to simplify. At this stage, we leave it to future efforts to implement more complicated formulations.

The powder-keg governing equations 6a-6d describe the activity of a neural field in the limit of a very large number of neurons per unit volume. For a finite number of neurons, the deterministic representation given by equation 6d may be interpreted as an ensemble average, i.e., an average over many repetitions of the same experiment. It is easy to argue, however, that a realistic representation of the firing rate (even in a deterministic form) should include some information of other elements of the stochastic nature of the firing process: for example, the variance of membrane fluctuations should play a major role in the effective values of threshold levels for firing.

it seems reasonable to assume that the firing rate depends crucially on two elements: 1) on the probability of a neuron to fire (related to the proximity of the state of a given neuron to the threshold, which involves, say the variance of the membrane fluctuations, but possibly other/all moments of the probability density); and 2) the distribution of internal kinetic energy over the neural population.

Denote by *u*(*t*) the subthreshold, mean membrane depolarization. Invoking an ergodicity argument, the internal kinetic energy *u* defined above may be regarded as a time average of *u*(*t*). Assuming that the subthreshold *u*(*t*) is a time-integral of the activity of ion channels, as a random walk. Even if the mean internal kinetic energy *u* is fixed, neurons may fire as a response to the random walk *u*(*t*). Moreover, qualitatively speaking, neurons with higher depolarization are more likely to fire. Let *P*(*u*) be the firing probability of a neuron with internal kinetic energy *u*. The observations above imply that *P*(*u*) is a monotonically increasing function, with *P*(0) = 0 and *P*(*U*) = 1. Denote by *p*(*u*) the probability of a neuron to fire in the unit of time. Because a neuron fires instantaneously when it reaches the threshold level *U, p*(*U*) = ∞.

As discussed in section 4, the distribution of *u* over the population of neurons in an element of volume is characterized by a probability density function *f*_u_(u), which may be written as

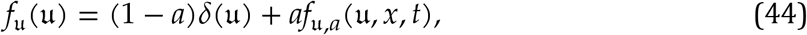

where the first term denotes the sub-population that is in refractory state (kinetic energy *u* = 0, where *δ* is the Dirac delta function), and *f*_u,*a*_(u, *x*, *t*) is the PDF component corresponding to active neurons. Taking into account the excitability *a*(*x*, *t*), the mean kinetic energy is

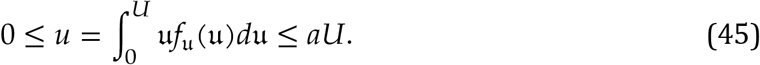

Because the firing rate as defined here is the number of firing events in the unit of time, the relationship between the firing rate of the population and its PDF *f*_u_(u) is given by the “activation functional”

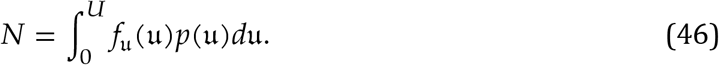

In the extreme case that *u* = *aU*, we have *f*_u,*a*_(u, *x*, *t*) = *δ*(u − *U*), thus firing rate *N* = ∞. As shown by equation 46, an accurate description of the time-evolution of the firing rate based on on the statistical state of the system involves fully detailed knowledge of the PDF *f*_u_(u). Alternatively, assuming that the moments of *f*_u_(u) completely characterize it, one could write

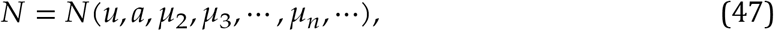

where *μ_n_* is the *n*-th moment of *f*_u_, where we assumed that the firing rate does not depend explicitly on time (note that the functional form *N* is different from the function *G* appearing in equation 5).

## A.1. Simplification of the activation function

Without further guidance about the the shape of *p*(u) and *f*_u_(u) (or all its moments), the only way to progress from equations 46 or 47 follows the beaten path of putting our hopes in assuming that the moments of *f*_u_ are well ordered at all times, i.e., *μ*_*n*+1_ ≪ *μ*_*n*_, and basically ignore all moments but the zeroth order (mean), i.e., write

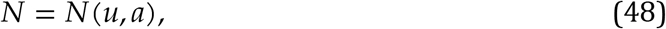

instead of equation 47. The simplified activation function should be a monotonically increasing function of *u* ∈ [0, *U*], with the end-point values and *N*(*u* = 0,*a*) = 0 and *N*(*u* = *U*,*a*) = ∞. A plausible functional form consistent with these constraints is

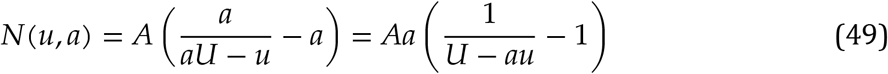

where the constant *A* is a measure of the intensity of endogenous membrane potential fluctuations. To further simplify the activation function, we may ignore the effect of *a* by setting for this calculation *a* ≈ 1 and effectively keeping only *u* as the controlling factor of the firing rate, which yields the expression given in equation 20, i.e.,

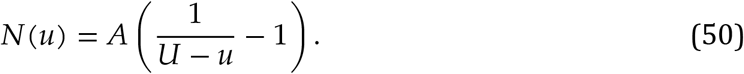

Equation 50 is arguably a “simplest” form of the activation function that describes the firing rate only as a function of the mean kinetic energy *u*. While this relation satisfies the leading order conditions stated above, it underestimates the firing rate in comparison with expression 49 but hopefully the difference is small unless *a* → 0, when a very large proportion of neurons are in absolute refractory state and *f*_u,*a*_(u, *x*, *t*) → *δ*(u − *U*). However, this condition implies that *f*_u_(u) is a U-shaped function, with large proportion of neurons in absolute refractory period, while the rest have a near threshold kinetic energy. This means the variance of *f*_u_(u) is relatively large. However, this cannot be a not a common condition of a network, because membrane depolarization in a network tends to be synchronized rather than the opposite (e.g., Wilson and Cowan, 1972a).

Therefore, we adopt for the activation function in this preliminary study the simple form 50, which is readily inverted to yield

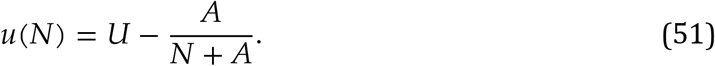

## A.2. The bounds of state variables *a* and *u*

A heuristic argument is as follows. From the definitions given in section 4, neurons in their absolute refractory period correspond to *u* = 0. Because *a* is the fraction of neuron population not in the absolute refractory state, and *U* is the maximum value of kinetic energy of individual neurons, the maximum value of *u* is *Ua*, hence *u* < *Ua*. The energy *u* cannot exceed *Ua* because the activation function 46 has the property that *N* → ∞ as *u* → *aU*, and in this case a large number of neurons drop to the level *u* = 0. This logic is also true for the simplified version of the activation function 49 that rely *N* on both *u* and *a*. However, whether the mathematical form of the model obeys this reasoning depends on the form of the activation function. From equations 7

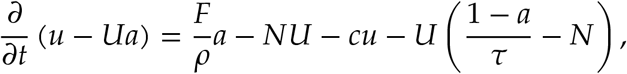

and using equation 1 obtains

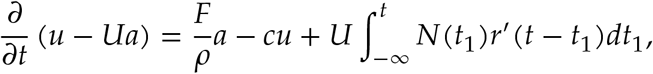

where 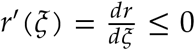, with input 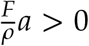, and the “inertial” terms 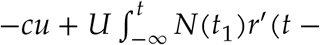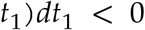. In the limit *N* → ∞ (equivalent to *u* → *Ua*), 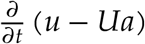 remains finite; in contrast, 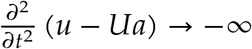, thus would not allow to exceed *Ua*. The derivative is negative if the forcing term 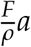 is negligible, but may become positive if the value of external input overwhelms the inertial (negative) part, which implies that *u* could exceed *Ua* (*a* decays as, *N* → ∞, and so does the contribution of the incoming of the forcing term 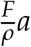). For the second derivative after some algebra, one obtains

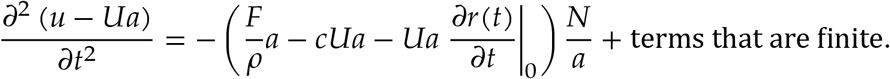

If 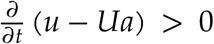 when *u* → *Ua*, i.e., if 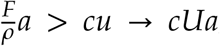, then 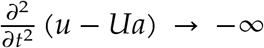, and consequently *u* would not exceed *Ua*. The activation function tells us that *N* approaches 0 when *u* approaches 0. The *u* < *Ua*. argument thus implies that *u* → 0 as *a* → 0. Moreover *N* → 0, as *u* → 0, thus, from equation 6c ∂*a*/∂*t* > 0, which insures that *a* cannot become negative.

Because the simplest form of the activation function 50 underestimates the firing rate, it is possible that it would indeed allow *u* to exceed the upper boundary *Ua* in extreme conditions when the external input *q* is very strong and the undervalued bursting rate is not large enough to cool down the system. However, assuming that the neural field operates far from this limiting case, the simple form 50 should provide a good approximation.

## APPENDIX B. Growth rate and phase lag for dual-type neural fields at equilibrium

Take *N*^*E*^ = *C*^*E*^*e*^*σt*^; 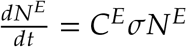 into Equation 39 with *α* = *E* we have

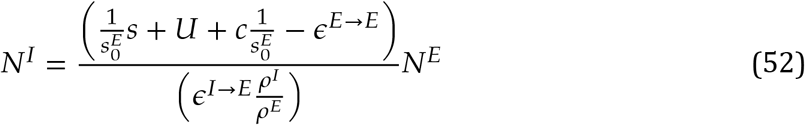

In which, the ratio of *N^I^* over *N^E^* is a complex number, phase of the ratio is the phase lag between inhibitory and excitatory populations.

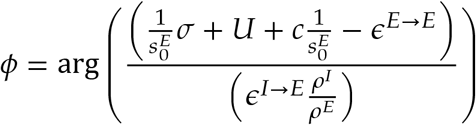

Take *N*^*I*^ = *C*^*I*^*e*^*σt*^; 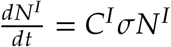 into Equation 39 with *α* = *I* we have

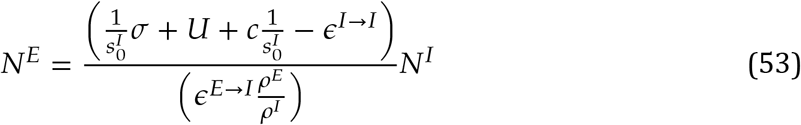

Combining Equation 52 with Equation 53 we know that.

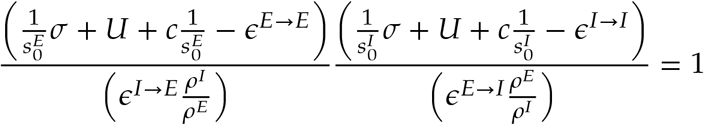

Then the complex oscillation frequency *σ* satisfy a quadratic equation

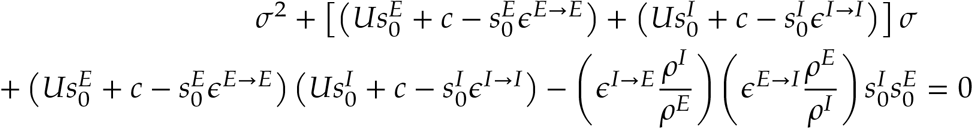

Thus the solutions of *σ* are

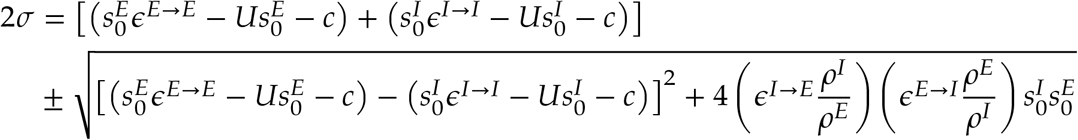

## APPENDIX C. Dispersion relation for dual-type neural fields

Take *N*^*E*^ = *C*^*E*^*e*^*i*(*kx*+*σt*)^; 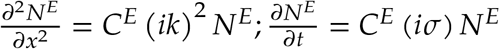 into Equation 38a with *α* = *E* we have.

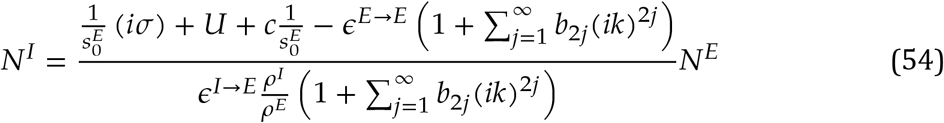

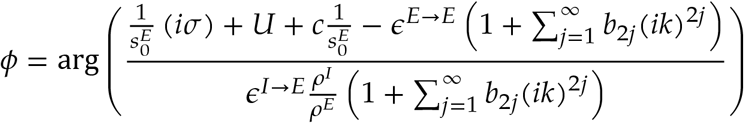

Take *N*^*I*^ = *C*^*I*^ *e*^*i*(*kx*+*σt*)^; 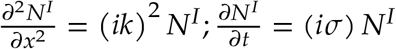 into Equation 38a with *α* = *I* we have.

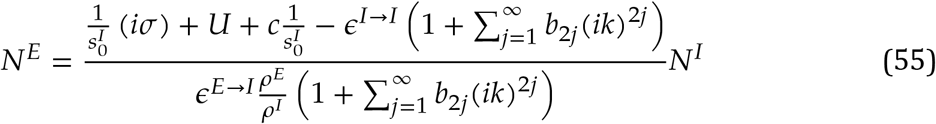

Combining Equation 54 with Equation 55 we know that.

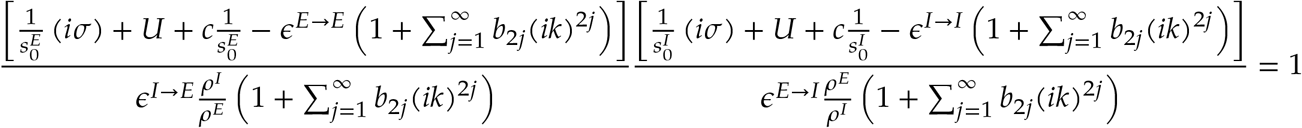

Then the complex oscillation frequency *σ* as a function of *k* satisfy a quadratic equation that

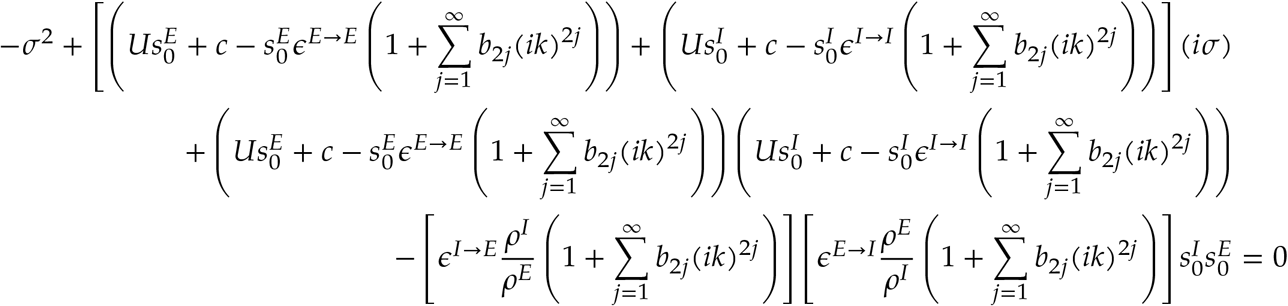

Thus the solutions of *σ* are

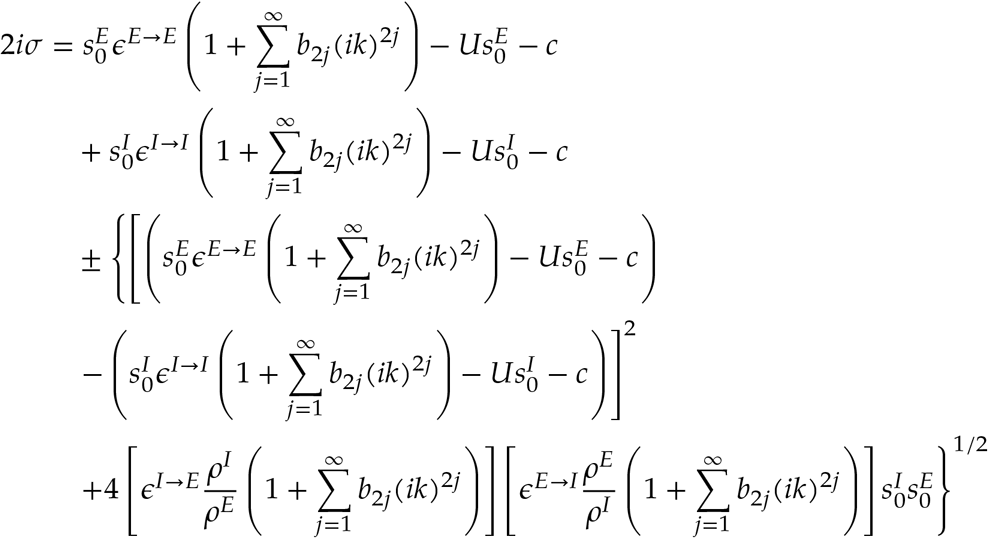

## Footnotes

1 The ideas below are elementary. We discuss them here only because they reflect a certain choice of terms, and for benefit of readers less familiar with statistical physics.

2 The “mesooscopic collective activity” concept is identical to Freeman’s [1975b] “mass action”. We prefer “collective action “ because the word “mass” has a reserved meaning in physics.

3 The ideas below are elementary. We discuss them here only because they reflect a certain choice of terms, and for benefit of readers less familiar with statistical physics.

4 A classical example of macroscopic behavior qualitatively distinct from microscopic physics is Boltzmann’s H-theorem for the idea gas. The (microscopic) dynamics of the gas particles is Hamiltonian, conservative and reversible; the (macroscopic) dynamics of the entire system is irreversible toward equilibrium (e.g., Boltzmann, 1872, 2003, Alexeev, 2004, Pathria and Beale, 2011).

5 This should be interpreted in the same sense as the statement “All cars on the road are Camrys”. The cars the all have the same mechanical characteristics, but can travel at different speeds, accelerations, etc.

6 These values are given for illustration purposes only; in actuality they depend on the type of neuron considered.

7 The term “potential” is used here in its literal sense, describing energy that is available, but not “realized”, or released.

8 The term volume is used in the general sense of measure, e.g., area for a two-dimensional network.

## References

B. V. Alexeev. Generalized Boltzmann Physical Kinetics. Elsevier, 2004.

P. G. Allen and F. S. Collins. Toward the final frontier: the human brain. The Wall Street Journal, 2013.

D.G. Amaral, H.E. Scharfman, and P. Lavenex. The dentate gyrus: fundamental neuroanatomical organization (dentate gyrus for dummies). Progress in Brain Research, 163:3–22, 2007. doi: 10.1016/S0079-6123(07)63001-5.

S. Amari. Homogeneous nets of neuron-like elements. Biological Cybernetics, 17:211–220, 1975.

S. Amari. Dynamics of pattern formation in lateral-inhibition type neural fields. Biological Cybernetics, 27, 1977.

S. Amari. Heaviside world: Excitation and self-organization of neural fields. Neural Fields: Theory and Applications, pages 97–118, 03 2014. doi: 10.1007/978-3-642-54593-1_3.

Bénédicte Amilhon, Carey YL Huh, Frédéric Manseau, Guillaume Ducharme, Heather Nichol, Antoine Adamantidis, and Sylvain Williams. Parvalbumin interneurons of hippocampus tune population activity at theta frequency. Neuron, 86(5):1277–1289, 2015.

R. Apps and M. Garwicz. Anatomical and physiological foundations of cerebellar information processing. Nature Reviews Neuroscience, 6.4:297, 2005. doi: 10.1038/nrn1646.

V. I. Arnold. Mathematical Methods of Classical Mechanics, volume 60 of Graduate Texts in Mathematics. Springer, 1974. ISBN 978-0387968902.

S.J. Aton, C. Broussard, M. Dumoulin, J. Seibt, A. Watson, T. Coleman, and M.G. Frank. Visual experience and subsequent sleep induce sequential plastic changes in putative inhibitory and excitatory cortical neurons. Proceedings of the National Academy of Sciences of the United States of America, 110:3101–3106, 2013.

A. Attardo, J. E. Fitzgerald, and M. J. Schnitzer. Impermanence of dendritic spines in live adult ca1 hippocampus. Nature, 523(7562):592–596, 2015. ISSN 1476-4687. doi: 10.1038/nature14467. URL https://www.ncbi.nlm.nih.gov/pubmed/26098371.

Per Bak, Chao Tang, and Kurt Wiesenfeld. Self-organized criticality. Physical Review A, 38:364–375, 1988.

M. Bartos, I. Vida, and P. Jonas. Synaptic mechanisms of synchronized gamma oscillations in inhibitory interneuron networks. Nature Reviews Neuroscience, 8:45–56, 2007.

John M. Beggs and Dietmar Plenz. Neuronal Avalanches in Neocortical Circuits. Journal of Neuroscience, 23(35):11167–11177, 2003.

John M. Beggs and Nicholas Timma. Being Critical of Criticality in the Brain. Frontiers in Physiology, 3:163, 2012.

R.W. Berg, A. Willumsen, and H. Lindén. When networks walk a fine line: balance of excitation and inhibition in spinal motor circuits. Current Opinion in Physiology, 8:76–83, 2019. ISSN 2468-8673.

R. L. Beurle and Bryan Harold Cabot Matthews. Properties of a mass of cells capable of regenerating pulses. Philosophical Transactions of the Royal Society of London. Series B, Biological Sciences, 240(669):55–94, 1956. doi: 10.1098/rstb.1956.0012. URL https://royalsocietypublishing.org/doi/abs/10.1098/rstb.1956.0012.

L. Boltzmann. Weitere Studien über das Wärmegleichgewicht unter Gasmolekülen. Sitzungsberichte Akademie der Wissenschaften, 66:275–370, 1872.

L. Boltzmann. History of Modern Physical Sciences: Volume 1, chapter Further Studies on the Thermal Equilibrium of Gas Molecules, pages 262–349. World Scientific, 2003.

C.A. Bosman, J.M. Schoffelen, N. Brunet, R. Oostenveld, A.M. Bastos, T. Womelsdorf, B. Rubehn, T. Stieglitz, P. De Weerd, and P. Fries. Attentional stimulus selection through selective synchronization between monkey visual areas. Neuron, 75(5):875–88, 2012.

A. Bragin, G. Jando, Z. Nadasdy, J. Hekte, K. Wise, and G. Buzsáki. Gamma (40-100 Hz) oscillation in the hippocampus of the behaving rat. Journal of Neuroscience, 15:47–60, 1995.

M. Breakspear and C.J. Stam. Dynamics of a neural system with a multiscale architecture. Phil. Trans. R. Soc. Lond. B, pages 1051–1074, 2005.

M. Breakspear, L.M. Williams, and C.J. Stam. A novel method for the topographic analysis of neural activity reveals formation and dissolution of “dynamic cell assemblies”. J. Comput. Neurosc, 16:49–68, 2004.

Michael Breakspear. Dynamic models of large-scale brain activity. Nature neuroscience, 20(3):340, 2017.

Michael Breakspear, John R Terry, and Karl J Friston. Modulation of excitatory synaptic coupling facilitates synchronization and complex dynamics in a biophysical model of neuronal dynamics. Network: Computation in Neural Systems, 14(4):703–732, 2003.

P.C. Bressloff. Spatiotemporal dynamics of continuum neural fields. Journal of Physics A: Mathematical and Theoretical, 45(3):033001, dec 2011. doi: 10.1088/1751-8113/45/3/033001.

G. Buzsáki. Theta oscillations in the hippocampus. Neuron, 33:325–340, 2002.

G. Buzsáki. Rhythms of the Brain. Oxford University Press, 2006.

G. Buzsaki. Rhythms of the Brain. Oxford University Press, 2006.

G. Buzsáki. Hippocampal sharp wave-ripple: A cognitive biomarker for episodic memory and planning. Hippocampus, 25(10):1073–1188, 2015.

G. Buzsáki and A. Draguhn. Neuronal oscillations in cortical networks. Science, 304.5679: 1926–1929, 2004.

D. Cai, L. Tao, M. Shelley, and D.W. McLaughlin. An effective kinetic representation of fluctuation-driven neuronal networks with application to simple and complex cells in visual cortex. Proceedings of the National Academy of Science USA, 101:7757–7562, 2004.

H. B. Callen. Thermodynamics. John Wiley & Sons, Inc., 1960.

Luísa Castro and Paulo Aguiar. Phase precession through acceleration of local theta rhythm: a biophysical model for the interaction between place cells and local inhibitory neurons. Journal of computational neuroscience, 33(1):141–150, 2012.

Stephen Coombes, Peter beim Graben, Roland Potthast, and James Wright, editors. Neural fields, Theory and appliocations. Springer, 2014.

Stephen; Gabriel J. Lord; Markus R. Owen. Coombes. Waves and bumps in neuronal networks with axo-dendritic synaptic interactions. Physica D: Nonlinear Phenomena, 178(3–4):219–241, 2003.

J. D. Cowan, J. Neuman, and W. van Drongelen. Wilson–Cowan Equations for Neocortical Dynamics. The Journal of Mathematical Neuroscience, 6(1):1:24, 2016.

Rodica Curtu and Bard Ermentrout. Oscillations in a refractory neural net. Journal of mathematical biology, 43(1):81–100, 2001.

FH Lopes Da Silva, A. Hoeks, H. Smits, and L. H. Zetterberg. Model of brain rhythmic activity, the alpha-rhythm of the thalamus. Kybernetik, 12(1):27–37, 1974.

G. Deco, V.K. Jirsa, P.A. Robinson, M. Breakspear, and K. Friston. The dynamic brain: From spiking neurons to neural masses and cortical fields. PLoS Computational Biology, 4(8): e1000092, 2008. doi: 10.1371/journal.pcbi.1000092.

Gustavo Deco, Viktor Jirsa, Anthony R McIntosh, Olaf Sporns, and Rolf Kötter. Key role of coupling, delay, and noise in resting brain fluctuations. Proceedings of the National Academy of Sciences, 106(25):10302–10307, 2009.

R. Desimone and J. Duncan. Neural Mechanisms of Selective Visual Attention. Annual Review of Neuroscience, 18:193–222, 1995.

K. Diba, A. Amarasingham, K. Mizuseki, and G. Buzsáki. Milliappond timescale synchrony among hippocampal neurons. J Neurosci, 34(45):14984–94, 2014. ISSN 1529-2401. doi: 10.1523/JNEUROSCI.1091-14.2014. URL http://www.ncbi.nlm.nih.gov/pubmed/25378164.

G. M. Edelman and J. A. Gally. Degeneracy and complexity in biological systems. Proc Natl Acad Sci U S A, 98(24):13763–8, 2001. ISSN 0027-8424. doi: 10.1073/pnas.231499798. URL https://www.ncbi.nlm.nih.gov/pubmed/11698650.

Gerald M Edelman. Neural Darwinism: The theory of neuronal group selection. Basic Books, 1987. ISBN 0465049346.

H. Eichenbaum. Barlow versus Hebb: When is it time to abandon the notion of feature detectors and adopt the cell assembly as the unit of cognition? Neurosci Letters, 2017.

Sami El Boustani and Alain Destexhe. A master equation formalism for macroscopic modeling of asynchronous irregular activity states. Neural computation, 21(1):46–100, 2009.

D.F. English, S. McKenzie, T. Evans, Kim K., Yoon E., and G. Buzsaki. Pyramidal cell-interneuron circuit architecture and dynamics in hippocampal networks. neuron, 96(2):505–520, 2017.

Bard Ermentrout. Neural networks as spatio-temporal pattern-forming systems. Reports on progress in physics, 61(4):353, 1998. ISSN 0034-4885.

G Bard Ermentrout and J Bryce McLeod. Existence and uniqueness of travelling waves for a neural network. Proceedings of the Royal Society of Edinburgh Apption A: Mathematics, 123(3):461–478, 1993.

G.B Ermentrout and D. Kleinfeld. Traveling electrical waves in cortex: insights from phase dynamics and speculation on a computational role. Neuron, 29:33–44, 2001.

Nicolas Fourcaud and Nicolas Brunel. Dynamics of the firing probability of noisy integrate- and-fire neurons. Neural computation, 14(9):2057–2110, 2002.

W. J. Freeman. Mass action in the nervous system. New York: Academic Press, 1975a.

W. J. Freeman. A proposed name for aperiodic brain activity: stochastic chaos. Neural Networks, 13:11–13, 2000a. doi: 10.1016/S0893-6080(99)00093-3.

W. J. Freeman. Neurodynamics: An exploration in mesoscopic brain dynamics. Perspectives in Neural Computing. Springer-Verlag London, 2000b.

W. J. Freeman. A cinematographic hypothesis of cortical dynamics in perception. International Journal of Psychophysiology, 60(2):149–161, 2006. doi: 10.1016/j.ijpsycho.2005.12.009.

W. J. Freeman and G. Vitiello. Nonlinear brain dynamics as macroscopic manifestation of underlying many-body field dynamics. Physics of Life Reviews, 3:93–118, 2006. doi: doi:10.1016/j.plrev.2006.02.001.

Walter J Freeman. Mass action in the nervous system: examination of the neurophysiological basis of adaptive behavior through the EEG. Academic Press New York:, 1975b. ISBN 0122671503.

Walter J. Freeman. Nonlinear gain mediating cortical stimulus-response relations. Biological Cybernetics, 33(4):237–247, 1979.

Walter J. Freeman. The physiology of perception. Scientific American, 264(2), 1991.

Walter J. Freeman and G. Vitiello. Vortices in brain waves. International Journal of Modern Physics B, 24(17):3269–3295, 2010.

P. Fries. A mechanism for cognitive dynamics: neuronal communication through neuronal coherence. Trends in Cognitive Sciences, 9:474–480, 2005.

S. Frisch. How cognitive neuroscience could be more biological—and what it might learn from clinical neuropsychology. Frontiers in Human Neuroscience, 8(541):1–13, 2014.

U. Frisch. Turbulence, The legacy of A.N Kolmogorov. Cambridge University Press, 1995.

K. Friston. The free-energy principle: a unified brain theory? Nat Rev Neurosci, 11(2):127–38, 2010. ISSN 1471-0048. doi: 10.1038/nrn2787. URL https://www.ncbi.nlm.nih.gov/pubmed/20068583.

L.Z. Garamszegi and M. Eens. The evolution of hippocampus volume and brain size in relation to food hoarding in birds. Ecology Letters, 7.12:1216–1224, 2004.

Wulfram Gerstner, Werner M. Kistler, Richard Naud, and Liam Paninski. Neuronal dynamics: From single neurons to networks and models of cognition. Cambridge University Press, 2014.

J. W. Gibbs. Elementary principles in statistical mechanics. Longmans, Green and Co., 1902.

H. Goldstein, C. P. Poole, and J. L. Safko. Classical Mechanics. Addison-Wesley, 2014. ISBN 978-0201657029.

J.D. Green and A.A. Arduini. Hippocampal electrical activity in arousal. Journal of Neurophysiology, 17:533–557, 1954.

J.D. Green and X. Machne. Unit activity of rabbit hippocampus. American Journal of Physiology, 181:219–224, 1955.

LM Harrison, O David, and KJ Friston. Stochastic models of neuronal dynamics. Philosophical Transactions of the Royal Society B: Biological Sciences, 360(1457):1075–1091, 2005.

K. Hasselmann. On the non-linear energy transfer in a gravity-wave spectrum part 1. general theory. Journal of Fluid Mechanics, 12(4):481–500, 1962. doi: 10.1017/S0022112062000373.

M.E. Hasselmo. f I had a million neurons: Potential tests of cortico-hippocampal theories. Progess in Brain Research, 219:1–19, 2015.

Ryoma Hattori, Kishore V Kuchibhotla, Robert C Froemke, and Takaki Komiyama. Functions and dysfunctions of neocortical inhibitory neuron subtypes. Nature neuroscience, 20(9): 1199, 2017.

D.O. Hebb. The organization of behavior: A neuropsychological theory. Wiley, New York, 1949.

A. L. Hodgkin and A. F. Huxley. A quantitative description of membrane current and its application to conduction and excitation in nerve. J Physiol, 117(4):500–44, 1952. ISSN 0022-3751. URL http://www.ncbi.nlm.nih.gov/pubmed/12991237.

A. J. Holtmaat, J. T. Trachtenberg, L. Wilbrecht, G. M. Shepherd, X. Zhang, G. W. Knott, and K. Svoboda. Transient and persistent dendritic spines in the neocortex in vivo. Neuron, 45(2):279–291, 2005. doi: 10.1016/j.neuron.2005.01.003.

Christopher J Honey, Rolf Kötter, Michael Breakspear, and Olaf Sporns. Network structure of cerebral cortex shapes functional connectivity on multiple time scales. Proceedings of the National Academy of Sciences, 104(24):10240–10245, 2007.

Rebecca Hoyle and Rebecca B. Hoyle. Pattern formation: an introduction to methods. Cambridge University Press, 2006.

X. Huang, W. C. Troy, Q. Yang, H. Ma, C. R. Laing, S. J. Schiff, and J. Y. Wu. Spiral waves in disinhibited mammalian neocortex. J Neurosci, 24(44):9897–902, 2004. ISSN 1529-2401. doi: 10.1523/JNEUROSCI.2705-04.2004. URL http://www.ncbi.nlm.nih.gov/pubmed/15525774.

Ben H. Jansen and Vincent G. Rit. Electroencephalogram and visual evoked potential generation in a mathematical model of coupled cortical columns. Biological cybernetics, 73(4): 357–366, 1995.

R.A. Jirsa, V.K. & Stefanescu. Neural population modes capture biologically realistic large scale network dynamics. Bull. Math. Biol, 73:325–343, 2011.

V. K. Jirsa and H. Haken. Field Theory of Electromagnetic Brain Activity. Physical Review Letters, 77(5):960–963, 1996.

V. K. Jirsa and H. Haken. A derivation of a macroscopic field theory of the brain from the quasimicroscopic neural dynamics. Physica D, 77(5):960–963, 1997.

Viktor Jirsa, Olaf Sporns, Michael Breakspear, Gustavo Deco, and Anthony Randal McIntosh. Towards the virtual brain: network modeling of the intact and the damaged brain. Archives italiennes de biologie, 148(3):189–205, 2010.

M. Kardar. Statistical Physics of Fields. Cambridge University Press, 2007a. ISBN 978-0-521-87341-3.

M. Kardar. Statistical Physics of Particles. Cambridge University Press, 2007b. ISBN 978-0-521-87342-0.

A. I. Khinchin. Mathematical foundations of statistical mechanics. Dover Publications Inc., 1949.

Charles Kittel. Elementary Statistical Physics. Wiley, 1958.

A.N. Kolmogorov. The local structure of turbulence in incompressible viscous fluid for very large Reynolds numbers. Proceedings: Mathematical and Physical Sciences: Turbulence and Stochastic Process: Kolmogorov’s Ideas 50 Years On (Jul. 8, 1991), 434(1890)(30):9–13, 1941.

N. Kopell, G. B. Ermentrout, M. A. Whittington, and R. D. Traub. Gamma rhythms and beta rhythms have different synchronization properties. Proc Natl Acad Sci U S A, 97(4):1867–72, 2000. ISSN 0027-8424. URL https://www.ncbi.nlm.nih.gov/pubmed/10677548.

N. Kopell, C. Borgers, D. Pervouchine, P. Malerba, and A. Tort. Gamma and Theta Rhythms in Biophysical Models of Hippocampal Circuits. Springer, 2010.

H. S. Kudrimoti, C. A. Barnes, and B. L. McNaughton. Reactivation of hippocampal cell assemblies: effects of behavioral state, experience, and eeg dynamics. J Neurosci, 19(10):4090–101, 1999. ISSN 1529-2401. URL http://www.ncbi.nlm.nih.gov/pubmed/10234037.

Y. Kuramoto. Self-entrainment of a population of coupled non-linear oscillators. In International Symposium on Mathematical Problems in Theoretical Physics, pages 420–422. Lecture Springer Notes in Physics, vol 39, 1975.

Raima Larter, Brent Speelman, and Robert M Worth. A coupled ordinary differential equation lattice model for the simulation of epileptic seizures. Chaos: An Interdisciplinary Journal of Nonlinear Science, 9(3):795–804, 1999.

K. S. Lashley. Visual mechanisms, volume 301, chapter The Problem of Cerebral Organization in Vision, pages 301–322. Oxford, England: Jacques Cattell, 1942.

K. S. Lashley. Cerebral organization and behavior. Research Publications - Association for Research in Nervous and Mental Disease, 36(1–4):14–18, 1958.

Karl S Lashley, KL Chow, and Josephine Semmes. An examination of the electrical field theory of cerebral integration. Psychological review, 58(2):123, 1951. ISSN 1939-1471.

Klaus Linkenkaer-Hansen, Vadim V Nikouline, J Matias Palva, and Risto J Ilmoniemi. Long-range temporal correlations and scaling behavior in human brain oscillations. Journal of Neuroscience, 21(4):1370–1377, 2001. ISSN 0270-6474.

J.E Lisman and M.A. Idiart. Storage of 7 +/− 2 short-term memories in oscillatory subcycles. Science, 267:1512–1515, 1995.

R. Lorente de No. Physiology of the nervous system, chapter Architectonics and structure of the cerebral cortex, pages 291–330. Oxford University Press, 1938.

E.V. Lubenov and A.G. Siapas. Hippocampal theta oscillations are travelling waves. Nature, (459):534–539, 2009.

S. J. Luck, L. Chelazzi, S. A. Hillyard, and R. Desimone. Neural mechanisms of spatial selective attention in areas v1, v2, and v4 of macaque visual cortex. Annual Review of Neuroscience, 77:24–42, 1997.

Brian N Lundstrom, Matthew H Higgs, William J Spain, and Adrienne L Fairhall. Fractional differentiation by neocortical pyramidal neurons. Nature neuroscience, 11(11):1335, 2008.

C. Ly and D. Tranchina. Critical analysis of a dimension reduction by a moment closure method in a population density approach to neural network modeling. Neural Computation, 19: 2032–2092, 2007.

Wei Ji Ma, Jeffrey M Beck, Peter E Latham, and Alexandre Pouget. Bayesian inference with probabilistic population codes. Nature neuroscience, 9(11):1432–1438, 2006.

C. J. Maley. Toward analog neural computation. Minds and Machines, 28:77–91, 2018.

V. Mante, D. Sussillo, K.V. Shenoy, and W.T. Newsome. Context-dependent computation by recurrent dynamics in prefrontal cortex. Nature, 503:78–84, 2013.

E. Marder and D. Bucher. Central pattern generators and the control of rhythmic movements. Current Biology, 11:R986–996, 2001.

André C. Marreiros, Jean Daunizeau, Stefan J. Kiebel, and Karl J. Friston. Population dynamics: variance and the sigmoid activation function. Neuroimage, 42(1):147–157, 2008.

B. L. McNaughton, C. A. Barnes, J.L. Gerrard, K. Gothard, M.W. Jung, J.J. Knierim, H. Kudrimoti, Y. Qin, W.E. Skaggs, M. Suster, and Weaver. Deciphering the hippocampal polyglot: the hippocampus as a path integration system. Journal of Experimental Biologuy, 199(1):173–185, Sep 1996. ISSN 1432-1106. doi: 10.1007/BF00237147. URL https://doi.org/10.1007/BF00237147.

M. Megias, Z. Emri, T.F. Freund, and A.I. Gulyas. Total number and distribution of inhibitory and excitatory synapses on hippocampal CA1 pyramidal cells. Neuroscience, 102(3):527–540, 2001.

Hil GE Meijer and Stephen Coombes. Travelling waves in a neural field model with refractoriness. Journal of mathematical biology, 68(5):1249–1268, 2014.

Jorge F Mejias, John D Murray, Henry Kennedy, and Xiao-Jing Wang. Feedforward and feedback frequency-dependent interactions in a large-scale laminar network of the primate cortex. Science advances, 2(11):e1601335, 2016.

P. Miller, C. D. Brody, R. Romo, and X. J. Wang. A recurrent network model of somatosensory parametric working memory in the prefrontal cortex. Cereb Cortex, 13(11):1208–18, 2003. ISSN 1047-3211. URL https://www.ncbi.nlm.nih.gov/pubmed/14576212.

John G Milton, Po Hsiang Chu, and Jack D Cowan. Spiral waves in integrate-and-fire neural networks. In Advances in neural information processing systems, pages 1001–1006, 1993.

L. Muller, F. Chavane, J. Reynolds, and T. J. Sejnowski. Cortical travelling waves: mechanisms and computational principles. Nat Rev Neurosci, 19(5):255–268, 2018a. ISSN 1471-0048. doi: 10.1038/nrn.2018.20. URL https://www.ncbi.nlm.nih.gov/pubmed/29563572.

Lyle Muller, Alexandre Reynaud, Frédéric Chavane, and Alain Destexhe. The stimulus-evoked population response in visual cortex of awake monkey is a propagating wave. Nature communications, 5(1):1–14, 2014.

Lyle Muller, Frederic Chavane, John Reynolds, and Terrence J. Sejnowski. Cortical travelling waves: mechanisms and computational principles. Nature Reviews Neuroscience, 19:255–268, 2018b.

S.V. Nazarenko. Wave Turbulence. Springer, 2011.

Garrett T Neske, Saundra L Patrick, and Barry W Connors. Contributions of diverse excitatory and inhibitory neurons to recurrent network activity in cerebral cortex. Journal of Neuroscience, 35(3):1089–1105, 2015.

A Newell. Lectures on Wave Turbulence and Intermittency, pages 227–271. Springer, 2002.

A.C. Newell, S.V. Nazarenko, and L. Biven. Wave turbulence and intermittency. Physica D: Nonlinear Phenomena, 152–153:520–550, 2001. doi: 10.1016/S0167-2789(01)00192-0.

Paul L Nunez and Ramesh Srinivasan. Electric fields of the brain: the neurophysics of EEG. Oxford University Press, USA, 2006. ISBN 019505038X.

P.L. Nunez. The brain wave equation: a model for eeg. Mathematical Bioscience, 21:279–297, 1974.

D. Nykamp and D. Tranchina. A population density method that facilitates large-scale modeling of neural networks: analysis and application to orientation tuning. Journal of Computational Neuroscience, 8:19–50, 2000.

Ahmet Omurtag, Bruce W. Knight, and Lawrence Sirovich. On the simulation of large populations of neurons. Journal of computational neuroscience, 8(1):51–63, 2000.

Remus Osan and Bard Ermentrout. Two dimensional synaptically generated traveling waves in a theta-neuron neural network. Neurocomputing, 38:789–795, 2001.

A. Parent and L.-N. Hazrati. Functional anatomy of the basal ganglia. i. the cortico-basal ganglia-thalamo-cortical loop. Brain Research Reviews, 20.1:91–127, 1995.

J. Patel, S. Fujisawa, A. Berenyi, S. Royer, and G. Buzsaki. Traveling Theta Waves along the Entire Septotemporal Axis of the Hippocampus. Neuron, 75(3):410–417, 2012.

J. Patel, E. W. Schomburg, A. Berenyi, S. Fujisawa, and G. Buzsaki. Local Generation and Propagation of Ripples along the Septotemporal Axis of the Hippocampus. Journal of Neuroscience, 33(43):17029–17041, 2013.

R. K. Pathria and P. D. Beale. Statistical mechanics. Elsevier, third edition, 2011.

H. Petsche and C. Stumpf. Topographic and toposcopic study of origin and spread of the regular synchronized arousal pattern in the rabbit. Electroencephalography and Clinical Neurophysiology, 12:589–600, 1960.

David J Pinto and G Bard Ermentrout. Spatially structured activity in synaptically coupled neuronal networks: I. traveling fronts and pulses. SIAM journal on Applied Mathematics, 62(1):206–225, 2001a. ISSN 0036-1399.

David J Pinto and G Bard Ermentrout. Spatially structured activity in synaptically coupled neuronal networks: Ii. lateral inhibition and standing pulses. SIAM Journal on Applied Mathematics, 62(1):226–243, 2001b.

D.J. Pinto, S.L. Patrick, W.C. Huang, and B.W. Connors. Initiation, propagation, and termination of epileptiform activity in rodent neocortex in vitro involve distinct mechanisms. Journal of Neuroscience, 25:8131–8140, 2005.

A.V. Rangan, G. Kovacic, and D. Cai. Kinetic theory for neuronal networks with fast and slow excitatory conductances driven by the same spike train. Physical Review E, 77:041915, 2008.

S. Ray and J. H. Maunsell. Different origins of gamma rhythm and highgamma activity in macaque visual cortex. PLoS Biol, 9(4):e1000610, 2011. ISSN 1545-7885. doi: 10.1371/journal.pbio.1000610. URL https://www.ncbi.nlm.nih.gov/pubmed/21532743.

J. H. Reynolds, L. Chelazzi, and R. Desimone. Competitive mechanisms subserve attention in macaque areas v2 and v4. Journal of Neuroscience, 19:1736–1753, 1999.

L.F. Richardson. Weather Prediction by Numerical Process. Cambridge Univ. Press, 1922.

P. A. Robinson, C. J. Rennie, and J. J. Wright. Propagation and stability of waves of electrical activity in the cerebral cortex. Physical Review E, 56(1):826–840, 1997.

PA Robinson. Patchy propagators, brain dynamics, and the generation of spatially structured gamma oscillations. Physical Review E, 73(4):041904, 2006.

PA Robinson, CJ Rennie, and DL Rowe. Dynamics of large-scale brain activity in normal arousal states and epileptic seizures. Physical Review E, 65(4):041924, 2002.

A. Sheremet, S. Burke, and A. Maurer. Movement enhances the nonlinearity of hippocampal theta. Journal of Neuroscience, 36(15):4218–4230, 2016a.

A. Sheremet, S. N. Burke, and A. P. Maurer. Movement enhances the non-linearity of hippocampal theta. J Neurosci, 36(15):4218–30, 2016b. ISSN 1529-2401. doi: 10.1523/JNEUROSCI.3564-15.2016. URL http://www.ncbi.nlm.nih.gov/pubmed/27076421.

A. Sheremet, J. P. Kennedy, Y. Qin, Y. Zhou, S. D. Lovett, S. N. Burke, and A. P. Maurer. Theta-gamma cascades and running speed. J Neurophysiol, 2018a. ISSN 1522-1598. doi: 10.1152/jn.00636.2018. URL https://www.ncbi.nlm.nih.gov/pubmed/30517044.

A. Sheremet, Y. Zhou, J.P. Kennedy, Y. Qin, S.N. Burke, and A.P. Maurer. Theta-gamma coupling: a nonlinear dynamical model. BioRXiv, doi: https://doi.org/10.1101/304238, 2018b.

A. Sheremet, J. P. Kennedy, Y. Qin, Y. Zhou, S.D. Lovett, Burke S. N., and A.P. Maurer. Thetagamma cascades and running speed. Journal of Neurophysiology, 121(2):444–458, 2019a. doi: 10.1152/jn.00636.2018.

Alex Sheremet, Yu Qin, Jack P Kennedy, Yuchen Zhou, and Andrew P Maurer. Wave turbulence and energy cascade in the hippocampus. Frontiers in Systems Neuroscience, 12:62, 2019b. ISSN 1662-5137. doi: doi: 10.3389/fnsys.2018.00062.

Alexandru Sheremet, Yu Qin, Jack P Kennedy, and Andrew Maurer. Mesoscale turbulence in the hippocampus. bioRxiv, page 217877, 2017.

Woodrow L Shew, Hongdian Yang, Shan Yu, Rajarshi Roy, and Dietmar Plenz. Information capacity and transmission are maximized in balanced cortical networks with neuronal avalanches. Journal of neuroscience, 31(1):55–63, 2011.

E. Stark, R. Eichler, L. Roux, S. Fujisawa, H. G. Rotstein, and G. Buzsáki. Inhibition-induced theta resonance in cortical circuits. Neuron, 80(5):1263–76, 2013. ISSN 1097-4199. doi: 10.1016/j.neuron.2013.09.033. URL http://www.ncbi.nlm.nih.gov/pubmed/24314731.

V.K. Stefanescu, R.A. & Jirsa. Reduced representations of heterogeneous mixed neural networks with synaptic coupling. Phys. Rev. E, 83, 2011.

S. H. Strogatz. From Kuramoto to Crawford: exploring the onset of synchronization in populations of coupled oscillators. Physica D, 143:1–20, 2000.

D. Sussillo and L.F. Abbott. Generating coherent patterns of activity from chaotic neural networks. Neuron, 63:544–557, 2009.

Edward Chace Tolman. The determiners of behavior at a choice point. Psychological Review, 45(1):1, 1938. ISSN 1939-1471.

David Tong. Kinetic Theory. Course Notes, Published online. 2012. URL https://www.damtp.cam.ac.uk/user/tong/kinetic.html.

R. D. Traub, N. Spruston, I. Soltesz, A. Kenneth, M. A. Whittington, and J. G. R. Jeffreys. Gamma-frequency oscillations: a neuronal population phenomenon, regulated by synaptic and intrinsic cellular processes, and inducing synaptic plasticity. Progress in Neurobiology, 55: 563–575, 1998.

A.J. Trevelyan, D. Sussillo, and R. Yuste. Feedforward inhibition contributes to the control of epileptiform propagation speed. Journal of Neuroscience, 2:3383–3387, 2007.

WC Troy. Wave phenomena in neuronal networks. Dissipative Solitons: From Optics to Biology and Medicine, pages 1–22, 2008.

C.H. Vanderwolf. Hippocampal electrical activity and voluntary movement in the rat. Electroencephalogarphy and Clinical Neurophysiology, 26:407–418, 1969.

X. J. Wang. Neurophysiological and computational principles of cortical rhythms in cognition. Physiol Rev, 90(3):1195–268, 2010. ISSN 1522-1210. doi: 10.1152/physrev.00035.2008. URL http://www.ncbi.nlm.nih.gov/pubmed/20664082.

J. A. White, C. C. Chow, J. Rit, C. Soto-Treviño, and N. Kopell. Synchronization and oscillatory dynamics in heterogeneous, mutually inhibited neurons. Journal of Computational Neuroscience, 5:5–16, 1998.

Gerald Beresford Whitham. Linear and nonlinear waves, volume 42. John Wiley & Sons, 2011. ISBN 1118031202.

M. A. Whittington, R. D. Traub, N. Kopell, B. Ermentrout, and E. H. Buhl. Inhibition-based rhythms: experimental and mathematical observations on network dynamics. Int J Psychophysiol, 38(3):315–36, 2000. ISSN 0167-8760. URL http://www.ncbi.nlm.nih.gov/pubmed/11102670.

H. R. Wilson and J. D. Cowan. Excitatory and inhibitory interactions in localized populations of model neurons. Biophys J, 12(1):1–24, 1972a. ISSN 0006-3495. doi: 10.1016/S0006-3495(72)86068-5. URL https://www.ncbi.nlm.nih.gov/pubmed/4332108.

H. R. Wilson and J. D. Cowan. Excitatory and inhibitory interactions in localized populations of model neurons. Biophysics Journal, 12:1–24, 1972b.

H. R. Wilson and J. D. Cowan. A mathematical theory of the functional dynamics of cortical and thalamic nervous tissue. Kybernetik, 13:55–80, 1973.

A.T. Winfree. The geometry of biological time. Springer Science and Business Media, 2001.

X.-J. Wong, K.-F. & Wang. A recurrent network mechanism of time integration in perceptual decisions. J. Neurosci., 26:1314–1328, 2006.

Mark W Woolrich and Klaas E Stephan. Biophysical network models and the human connectome. Neuroimage, 80:330–338, 2013.

J. J. Wright and D. T. Liley. Simulation of electrocortical waves. Biol Cybern, 72(4):347–56, 1995a. ISSN 0340-1200. URL https://www.ncbi.nlm.nih.gov/pubmed/7748961.

J. J. Wright and D. T. J. Liley. Simulation of electrocortical waves. Biological Cybernetics, 72(4):347–356, 1995b.

JJ Wright and DTJ Liley. Dynamics of the brain at global and microscopic scales: Neural networks and the eeg. Behavioral and Brain Sciences, 19(2):285–295, 1996.

JJ Wright, AA Sergejew, and DTJ Liley. Computer simulation of electrocortical activity at millimetric scale. Electroencephalography and clinical Neurophysiology, 90(5):365–375, 1994.

J. Y. Wu, Xiaoying Huang, and Chuan Zhang. Propagating waves of activity in the neocortex: what they are, what they do. Neuroscientist, 14(5):487–502, 2008. ISSN 1073-8584. doi: 10.1177/1073858408317066. URL http://www.ncbi.nlm.nih.gov/pubmed/18997124.

Peer Wulff, Alexey A Ponomarenko, Marlene Bartos, Tatiana M Korotkova, Elke C Fuchs, Florian Bähner, Martin Both, Adriano BL Tort, Nancy J Kopell, William Wisden, et al. Hippocampal theta rhythm and its coupling with gamma oscillations require fast inhibition onto parvalbumin-positive interneurons. Proceedings of the National Academy of Sciences, 106(9):3561–3566, 2009.

Z. Xiao, P. Y. Deng, C. Yang, and S. Lei. Modulation of gabaergic transmission by muscarinic receptors in the entorhinal cortex of juvenile rats. J Neurophysiol, 102(2):659–69, 2009. ISSN 0022-3077. doi: 10.1152/jn.00226.2009. URL http://www.ncbi.nlm.nih.gov/pubmed/19494196.

T. Xu, X. Yu, A. J. Perlik, W. F. Tobin, J. A. Zweig, K. Tennant, T. Jones, and Y. Zuo. Rapid formation and selective stabilization of synapses for enduring motor memories. Nature, 462(7275):915–9, 2009. ISSN 1476-4687. doi: 10.1038/nature08389. URL https://www.ncbi.nlm.nih.gov/pubmed/19946267.

V.E. Zakharov. Statistical theory of gravity and capillary waves on the surface of a finite-depth fluid. European Journal of Mechanics, B/Fluids, 18(3):327–344, 1999.

V.E. Zakharov, V.S. L’vov, and G. Falkcovich. Kolmogorov spectra of turbulence I. Springer Series in Nonlinear Dynamics. Springer-Verlag, 1992a.

V.E. Zakharov, V.S. L’Vov, and G. Falkovich. Kolmogorov spectra of turbulence 1. wave turbulence. Kolmogorov spectra of turbulence 1. Wave turbulence., by Zakharov, VE; L’vov, VS; Falkovich, G.. Springer, Berlin (Germany), 1992, 275 p., ISBN 3-540-54533-6, 1, 1992b.

Y. Zhou, A. Sheremet, Y. Qin, J. P. Kennedy, N. M. DiCola, S. N. Burke, and A. P. Maurer. Methodological considerations on the use of different spectral decomposition algorithms to study hippocampal rhythms. eNeuro, 6(4), 2019. ISSN 2373-2822. doi: 10.1523/ENEURO.0142-19.2019. URL https://www.ncbi.nlm.nih.gov/pubmed/31324673.

